# miRNA-Mediated Regulation of Gene Expression During Early Activation in Jurkat Cells

**DOI:** 10.1101/2025.06.15.659805

**Authors:** Pooja Mukherjee, Thyago Leal-Calvo, Lucas Ferguson, Jamie H. D. Cate

## Abstract

**Background:** T cell activation induces substantial changes in gene expression by rapidly increasing transcription and translation. Additionally, microRNAs play a crucial role in regulating protein expression in T cell physiology, adding a layer of complexity by fine-tuning protein levels. While various miRNAs have been implicated in T cell function, a systematic analysis of differentially expressed miRNAs during early T cell activation and identification of their mRNA targets remains mostly unknown.

**Results:** We investigated dynamic changes in global gene expression during early T cell activation using a multi-omics approach combining small RNA-seq, mRNA-seq and ribosome profiling. Our results show that most differential expression changes occur by 5 hours post-activation, with translational upregulation predominating over downregulation. From 5 to 12 hours, we observed modest transcriptional and translational reprogramming. We identified 9 miRNAs that are differentially expressed (DE) during early activation, with most changes occurring as early as 5 hours. We calculated translation efficiency (TE) and classified genes based on changes in both mRNA abundance and ribosome-protected fragments (RPFs). By integrating TE and miRNA expression data, we examined the relationship between TE group-specific regulation patterns and the number of miRNA binding sites. Interestingly, rather than observing a uniform downregulation of targets with 4 or more predicted DE miRNA binding sites, we found distinct regulatory patterns that varied with both activation time point and TE category.

**Conclusions:** Our data provide new insights into how genes associated with key events in T cell activation such as translation, cell proliferation, and immune signaling are regulated at both the transcriptional and translational levels. The observation that most regulatory changes occur within the first 5 hours post-activation highlights the rapid and coordinated nature of T cell responses. The differential patterns of target regulation, based on translation efficiency groups and miRNA binding site density, suggest a context-dependent role for miRNAs in shaping protein output. Future experiments will be required to functionally validate specific miRNA–target interactions and to explore their relevance in primary T cells *in vivo*. This study also lays the groundwork for identifying miRNA-based regulatory circuits for therapeutic modulation of T cell activity.

## Background

T cell activation results in a rapid increase in transcription and translation that can sustain the activated state for days [1–3]. T cell activation can be divided into “early” and “late” stages, with the “early” phase describing changes occurring within 4-6 hours of activation and the “late” phase encompassing changes 24 hours or later post-activation [4,5]. This process leads to a comprehensive remodeling of the cellular transcriptome and proteome, resulting in significant alterations in multiple pathways related to cellular metabolism and protein synthesis [2,6–8]. The main events following T cell activation include activation of cell surface markers, cytokine secretion, and cell proliferation. Various cell surface markers are upregulated at different stages of activation. For instance, CD69 appears early on T cell surfaces after stimulation but decreases later [9,10] and CD25, part of the IL-2 receptor, is notably high in activated T cells [11].

T cell activation greatly increases translation rates, starting around 60,000 proteins per minute in quiescent cells, surging to approximately 300,000 proteins per minute within the initial 6 hours, and peaking at about 800,000 proteins per minute by the 24-hour mark post-activation. This heightened synthesis persists for 1-2 days, priming the cells for rapid division [2,3]. While proteome and mRNA changes differ more during early activation, they become almost equal during the proliferation phase, a phase when activated T cells rapidly divide and expand in number [3,4,12,13]. Additionally, the decrease in differentially expressed (DE) proteins during late activation indicates convergence between CD4+ and CD8+ T cells in their proliferative stages, not evident at the mRNA level [4].

MicroRNAs (miRNAs) play a significant role in regulating protein expression within T cells. While a global downregulation of miRNA activity occurs late in T cell activation [14], some specific miRNAs are upregulated in both CD8+ and CD4+ activated cells [15,16]. Moreover, a distinct set of miRNAs may show varied expression patterns during the initial stages of activation, an aspect that has been minimally explored [17]. A considerable amount of variability has been observed in previous studies on T cell-specific miRNAs and their expression, which may be due to several factors, including the T cell subtype and origin, the methodology of *in vitro* stimulation, and the choice of miRNA expression assessment tools [15]. miRNAs, together with protein Argonaute 2, form a miRNA-induced silencing complex (miRISC) and primarily repress their targets through mRNA degradation and/or translation repression [18–20]. However, miRNAs can also positively regulate gene expression. For example, some miRNAs localize to the nucleus and activate transcription by binding to promoters [21–23], while others enhance translation by targeting non-seed sequences like AU-rich elements [24,25] or the 5’ terminal oligopyrimidine tract (5’ TOP) motif in gene 5’-untranslated regions (5’-UTRs) [26]. Some miRNAs can also act as decoys, binding to ribonucleoproteins in a seed-independent manner, thereby preventing their interaction with target mRNAs and thereby indirectly activating target mRNA translation [27]. Moreover, recent findings challenge the traditional seed-centric model of miRNA targeting by showing that extensive 3′ pairing alone, without seed complementarity, can drive functional repression, as demonstrated for miR-155 and miR-124 *in vitro* and in cells [28].

In cultured T cells, activation can be induced by engaging the T cell receptor (TCR) and a coactivating receptor such as CD28 using agonist anti-CD3 and anti-CD28 antibodies, or co-culturing T cells with antigen-presenting cells [29–32]. To elucidate the overarching mechanism of miRNA function in T cell activation, we analyzed global changes in miRNA and mRNA expression and translation in Jurkat cells activated for 5 hours (early activation stage) and 12 hours (mid activation stage) after treatment with anti-CD3 and anti-CD28 bead-bound antibodies. We performed small RNA-seq to examine miRNA dynamics and mRNA-seq combined with ribosome profiling to study transcription and translation dynamics (Fig 1a). We observed that transcriptional and translational regulation of gene expression from 0 hours to 5 hours affected different pathways than those observed comparing 5 hours to 12 hours after activation. For example, most T cell specific pathway genes are both transcriptionally and translationally upregulated at 5 hours, whereas these genes are not substantially differentially regulated between 5 and 12 hours. We also observed that most miRNA expression changes occurred by 5 hours, with minimal differences between 5 and 12 hours. We identified experimentally validated targets of the differentially expressed (DE) miRNAs using multiple miRNA target databases. Six out of the 9 DE miRNAs at 5 hours had targets predicted for genes in immune-related pathways. We also found that genes targeted by multiple DE miRNAs (≥4) exhibited distinct regulatory patterns involving both upregulation and repression. Our findings suggest that DE miRNAs may fine-tune gene expression by supporting an initial burst of translation during early activation, particularly for genes critical to proliferation, followed by a shift toward repression as T cells progress through activation. However, further studies will be needed to determine whether these effects are direct and mechanistically linked to the identified miRNAs.

**Figure 1.**
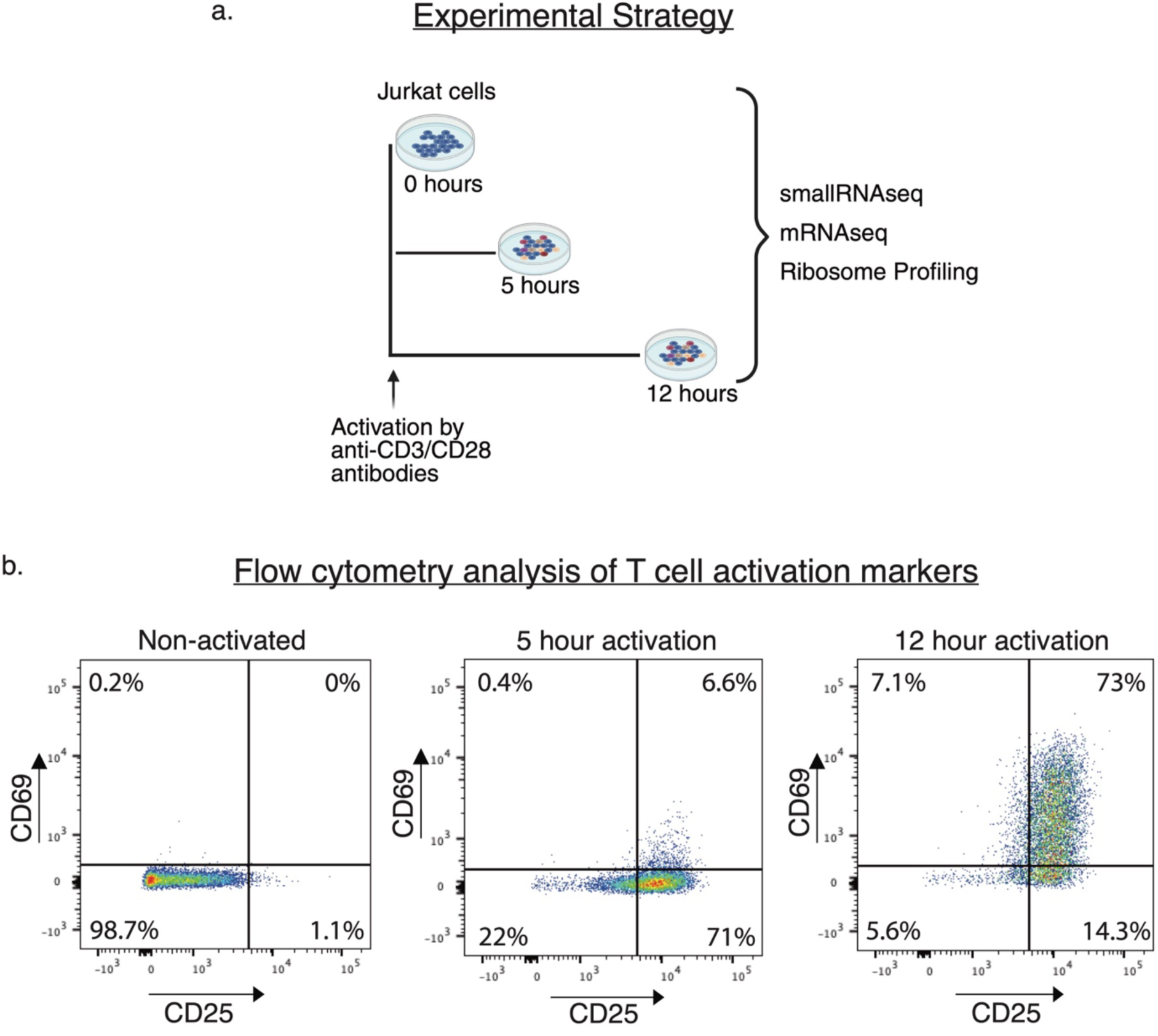
Experimental Strategy and Activation Validation of Jurkat cells. (a) Jurkat cells were stimulated with anti-CD3/CD28 antibodies, and samples were collected at 0, 5, and 12 hours post-activation. Small RNA sequencing (smallRNAseq), mRNA sequencing (mRNAseq), and ribosome profiling were performed to analyze miRNA expression, transcriptome changes, and translational regulation over time. This figure was created in https://BioRender.com. (b) Flow cytometry analysis of CD69 and CD25 surface markers in Jurkat cells before activation (non-activated) and after activation for 5 and 12 hours. The quadrants show the proportion of cells expressing CD69 and/or CD25 at each time point, with increased expression indicating T cell activation status.

## Results

### T cell specific genes are primarily upregulated at the transcriptional level during early activation

Previous studies revealed a burst in transcription and translation upon T cell activation [1–3]. While numerous studies have focused on identifying gene expression changes during late T cell activation (i.e. 24 hours or later post-activation), only a few have studied early activation [1,4,17]. We therefore examined the transcriptional and translational changes that occur during early to mid-stage activation of Jurkat cells (Fig 1a). We confirmed proper activation of Jurkat cells by assessing the expression of CD69 and CD25 surface markers at both 5 hours and 12 hours post-activation (Fig 1b). We then used these time points to perform mRNA deep sequencing and ribosome profiling under the same conditions to comprehensively understand the global mRNA expression changes following 5 hours and 12 hours of activation (Fig 1a, Suppl Fig 1a and Methods). For ribosome profiling, we used the newly-developed ordered two-template relay (OTTR) method, which first generates ribosome footprints using P1 nuclease, followed by a single-tube protocol for sequencing library preparation that integrates adaptors through reverse transcription [33]. We confirmed the optimal fragment length after library preparation (Suppl Fig 1b and Methods), and observed that 15-20% of the total reads in each sample represented putative ribosome protected mRNA fragments (Suppl Fig 1c). Using these data, we determined changes in transcript abundance and also calculated translation efficiency (TE), which is the ratio of ribosome-protected fragments (RPFs) within a gene’s coding sequence (CDS) to mRNA counts. TE represents how well each mRNA is translated in the cell [34].

Principal component analysis (PCA) of both mRNA-seq and ribosome profiling data revealed that the 5-hour samples are more closely correlated with the 12-hour samples than with the 0-hour samples, indicating early transcriptomic and translational shifts during T cell activation (Suppl Fig 2a). We also observed a significant proportion of genes differentially expressed at the 5 hour time point compared to no activation, with nearly the same number of genes upregulated and downregulated at the transcript level (Fig 2a, Suppl Fig 2b). Interestingly, two times as many genes are upregulated at the translational level compared to downregulated at 5 hours (Fig 2b and Suppl Fig 2b). The differences between 12 hours and 5 hours are less extensive overall (Fig 2c-d, Suppl Fig 2), but still show a pattern of more genes translationally upregulated compared to downregulated (Fig 2d and Suppl Fig 2b). Taken together, the global analysis of differential gene expression at 5 hours and 12 hours post-activation reveals most changes occur by 5 hours, with more genes translationally upregulated than downregulated, followed by more modest transcriptional and translational reprogramming occurring from 5 hours to 12 hours (Suppl. Fig 2). Notably, these effects are masked if comparisons are made just between 12 hours and 0 hours (Suppl. Fig 3), highlighting the importance of examining early time points after activation (here, 5 hours) (Suppl Tables S1-S3).

**Figure 2.**
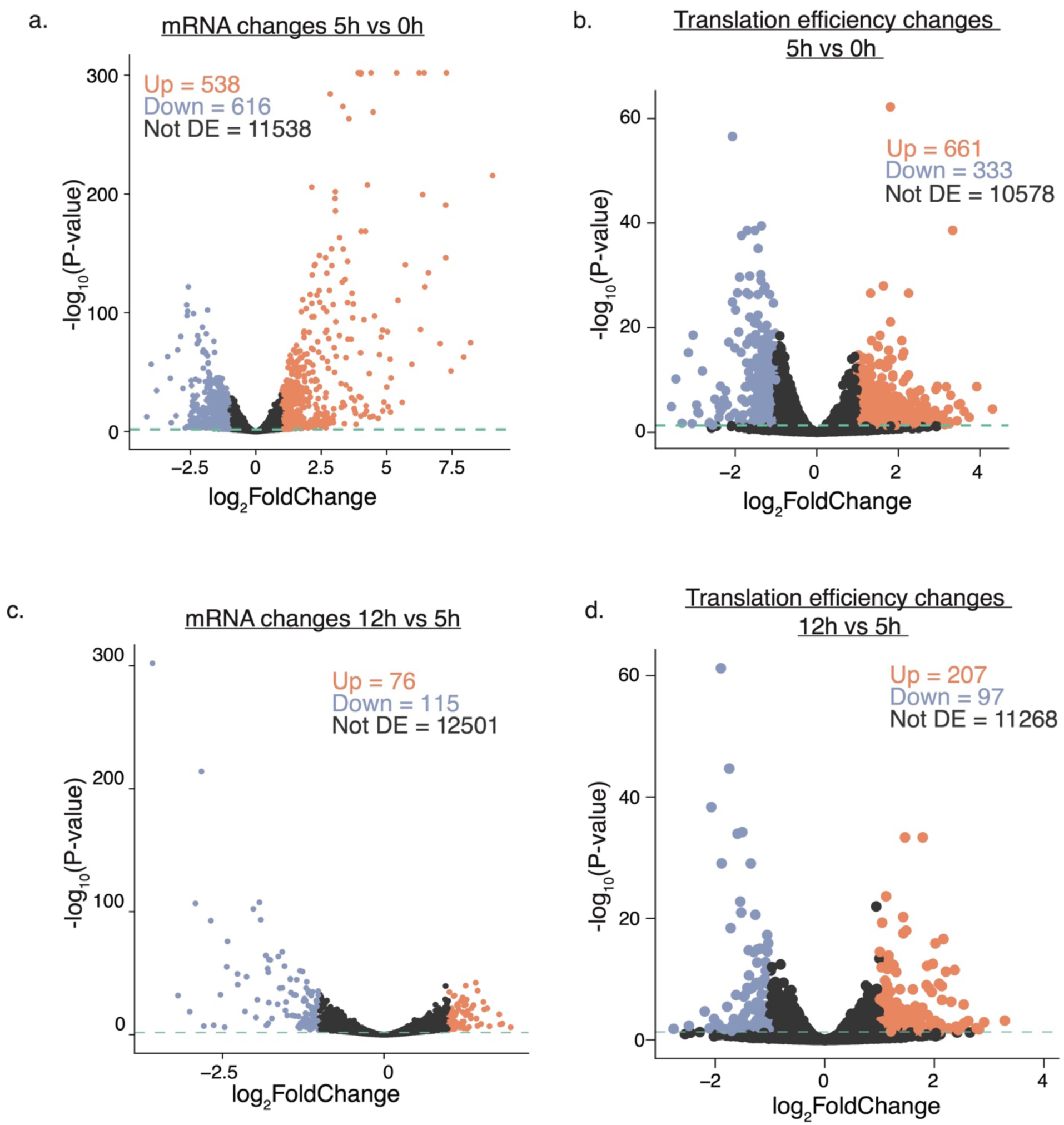
mRNA and Translation Efficiency Changes at 5 and 12 hours. (a-b) Volcano plots depicting mRNA expression (a) and translation efficiency (b) change at 5 hours post-activation in Jurkat cells. (c-d) Volcano plots depicting mRNA expression (c) translation efficiency (d) changes at 12 hours compared to 5 hours post-activation in Jurkat cells. Each dot represents a gene, with the x-axis showing log2 fold changes and the y-axis displaying -log10 p-values. Genes with significant upregulation are colored orange, and downregulated genes are in blue. The threshold used was FDR (Benjamini–Hochberg (BH) correction) ≤ 5% and |log2FC| ≥ 1.

Among the differentially expressed genes, we identified T cell activation specific genes that are transcriptionally regulated (Fig 3a, 3b) and translationally regulated (Fig 3c, 3d) at both time points. These include early response genes (e.g., *CD69*, *ZFP36L1*), transcription factors (e.g., *TBX21*, *STAT3*), and cell proliferation genes (e.g., *CCND3*, *KIT*). This pattern reflects coordinated transcriptional and translational activation of T cell functional programs. Notably, the early activation marker *CD69* was highly upregulated transcriptionally and also showed enhanced translation efficiency, suggesting tight post-transcriptional regulation at 5 hours (Fig. 3a, 3c). However, by 12 hours, *CD69* is downregulated transcriptionally (Fig 3b), preceding CD69 disappearance from the cell surface (Fig. 1b). A second activation marker in immune cells, *CD83*, shows a slightly different pattern of expression, with *CD83* upregulated transcriptionally and translationally at 5 hours (Fig. 3a, 3c). At 12 hours, both *CD83* transcript levels and TE drop in tandem (Fig. 3b, 3d).

**Figure 3.**
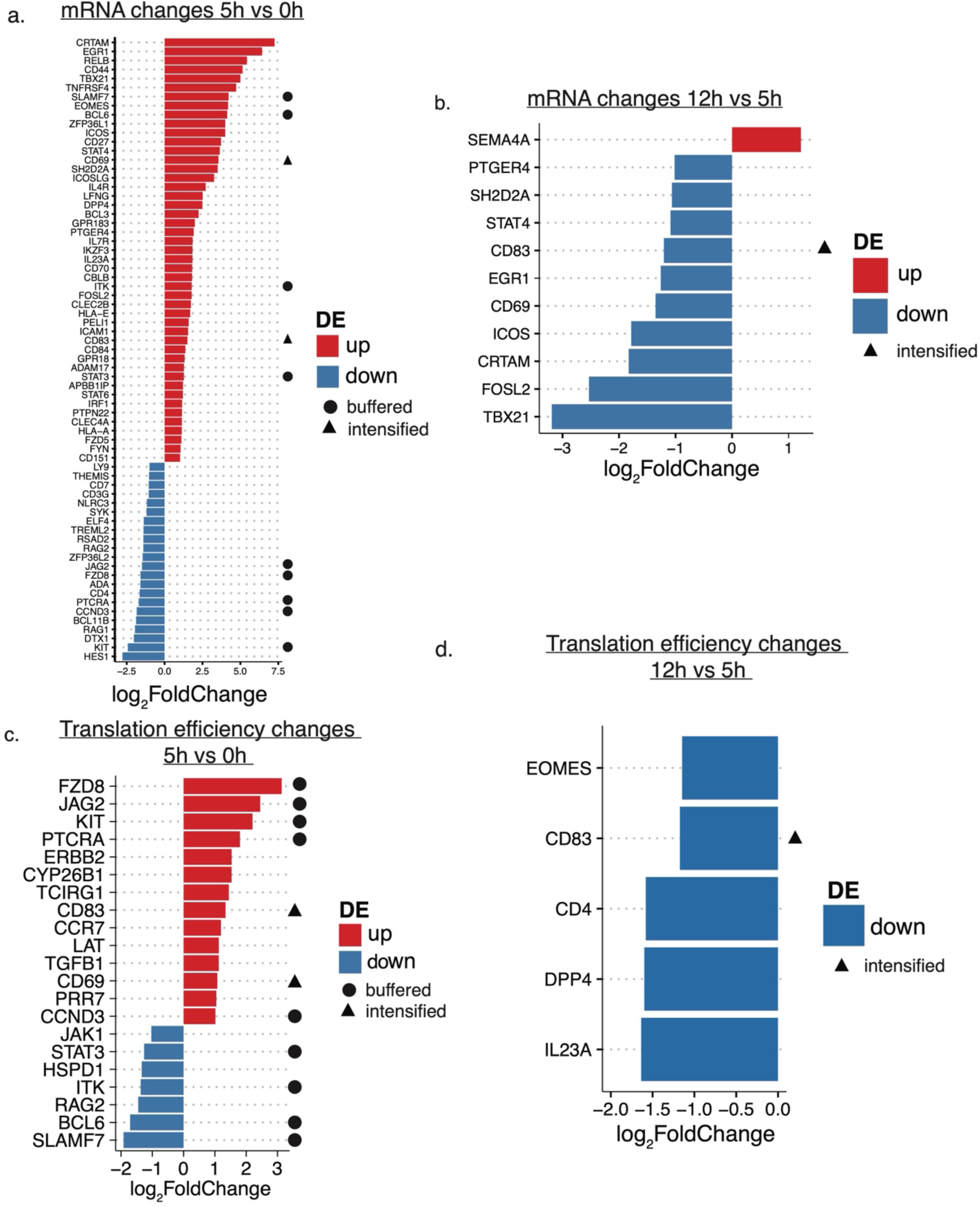
Differential expression of T cell activation-specific genes at different time points. (a–b) Bar plots showing upregulated and downregulated T cell activation-specific genes at the mRNA level at 5 hours (a) and 12 hours (b). (c–d) Bar plots showing changes in translation efficiency for T cell activation-specific genes at 5 hours (c) and 12 hours (d). The x-axis indicates gene names; the y-axis represents log2 fold changes. Triangles and circles indicate genes classified as intensified or buffered, respectively, as defined in Supplementary Figure 3. The threshold used was FDR (BH) ≤ 5% and |log2FC| ≥ 1.

The distinct patterns of *CD69* and *CD83* expression highlight the complex interplay of transcriptional and translational regulation observed in activated Jurkat cells at 5 hours and 12 hours. To more systematically analyze these patterns, we further classified genes which we could quantify at the translational level into four groups [34]: forwarded (both mRNA and RPF levels change in the same direction), buffered (mRNA levels and RPF levels change in the opposite direction, thereby buffering gene expression), exclusive (mRNA levels are unchanged but RPF changes) and intensified (mRNA and RPF levels change in the same direction, but with RPF changes more pronounced, thereby intensifying gene expression) [34] (Suppl Fig 4). This classification allows us to examine the patterns of mRNA and TE changes for each gene during T cell activation. Grouping of genes based on these patterns, followed by gene ontology (GO) analysis, revealed striking patterns of regulation of gene expression that changed from 0 hours to 5 hours and then from 5 hours to 12 hours (Fig 4 and Suppl Fig 5). GO over-representation analysis (ORA) showed that most of the forwarded gene categories at 5 hours are T cell or immune specific, meaning the increase in expression is mainly driven by transcription rather than translation, since both the mRNA and ribosome footprint levels change to similar extents (Fig 4 and Suppl Fig 5). However, by 12 hours, the changes in immune-related pathways become intensified (Fig 4), where changes in translation amplify changes in mRNA levels. By contrast, several cell division-specific gene categories fell into the intensified class at 5 hours, but reverted to the forwarded class by 12 hours (Fig. 4 and Suppl Fig 5). We also observed pathways associated with ribosome biogenesis regulated at the translational level (the exclusive category) at 5 hours, and also in the buffered category at both 5 hours and 12 hours (Suppl Fig 5). Thus, classifying genes based on changes in mRNA and ribosome footprints provides a better understanding of how genes associated with key events in T cell activation, such as translation and cell proliferation, are regulated. Furthermore, measuring changes in mRNA levels and RPFs at 5 hours revealed changes in gene expression regulation that would be missed with just measurements at 12 hours (Suppl Fig 6).

**Figure 4.**
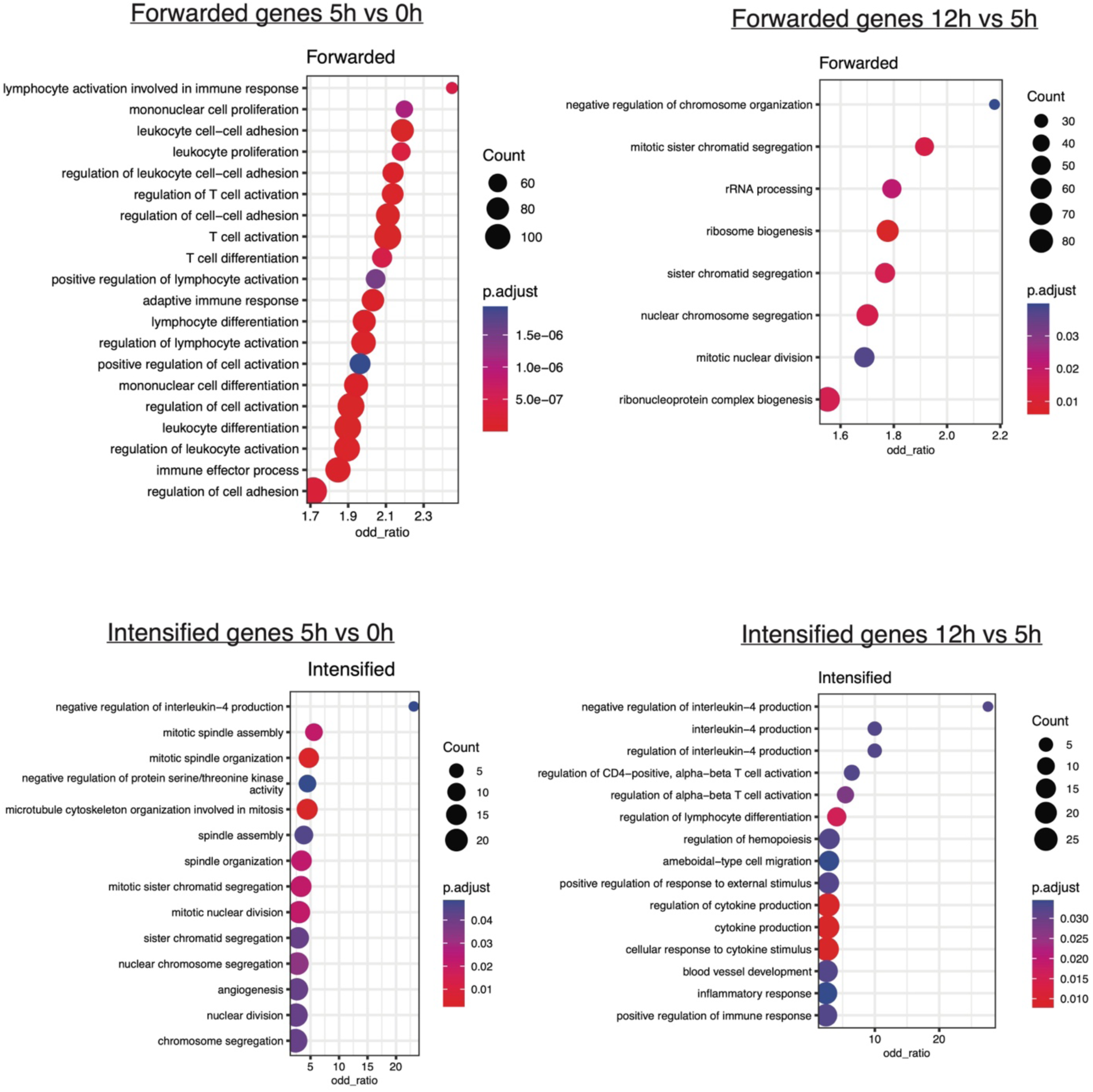
GO term enrichment analysis of forwarded and intensified genes at different time points. GO term enrichment analysis of forwarded and intensified genes at 5 hours and 12 hours. Dot plots display the enriched biological processes with the adjusted p-values (p.adjust) represented by the color scale. Dot size indicates the number of genes associated with each process. The threshold used was FDR (BH) ≤ 5%.

### Changes in miRNA Expression Occur During Early T Cell Activation

Next, we sought to understand miRNA expression changes during early and mid stages of T cell activation (Suppl Table S4 and S5). We detected a total of 636 miRNAs expressed (> 0 counts) by small RNA-seq, with 430 detected across all time points (Fig. 5a). We used a minimal cutoff of RNA counts > 0 across all samples to ensure that all detected miRNAs were included in our analysis. Notably, applying a more stringent cutoff of >10 counts per miRNA would exclude only one differentially expressed miRNA. Our data shows that miRNA expression in cells activated for 5 or 12 hours clustered with each other, distinct from the non-activated cells (0 hour time point), as determined by PCA (Fig. 5b). The relative expression levels of most miRNAs remained nearly constant across time points (Fig. 5c). However, among the observed miRNAs, 5 had increased expression at 5 hours post-activation (miR-132-3p, miR-132-5p, miR-146a-5p, miR-21-5p, miR-21-3p), and remained elevated at 12 hours post-activation (Suppl Fig 7, Suppl Table S4, Suppl Table S5). One decreased at 5 hours (miR-4521), and stayed repressed at 12 hours post-activation (Suppl Fig 7, Suppl Table S4, Suppl Table S5). At each timepoint, 3 additional miRNAs were differentially expressed when compared to non-activated cells (adj. P-value < 0.05, |Fold Change| ≥ 1.5) (Fig 5c, Suppl Table S4, Suppl Table S5). Many miRNAs generate two mature strands from their precursor molecules: the 5p arm from the 5’ end and the 3p arm from the 3’ end [35–37]. Both arms can be functional and can play distinct roles in gene regulation depending on specific conditions and cell types, adding complexity to miRNA-mediated processes [38–42]. Our data showed differential expression of both the 5p and 3p strands for miRNA-132 and miRNA-21 at both 5 hours and 12 hours (Suppl Table S4, Suppl Table S5).

**Figure 5.**
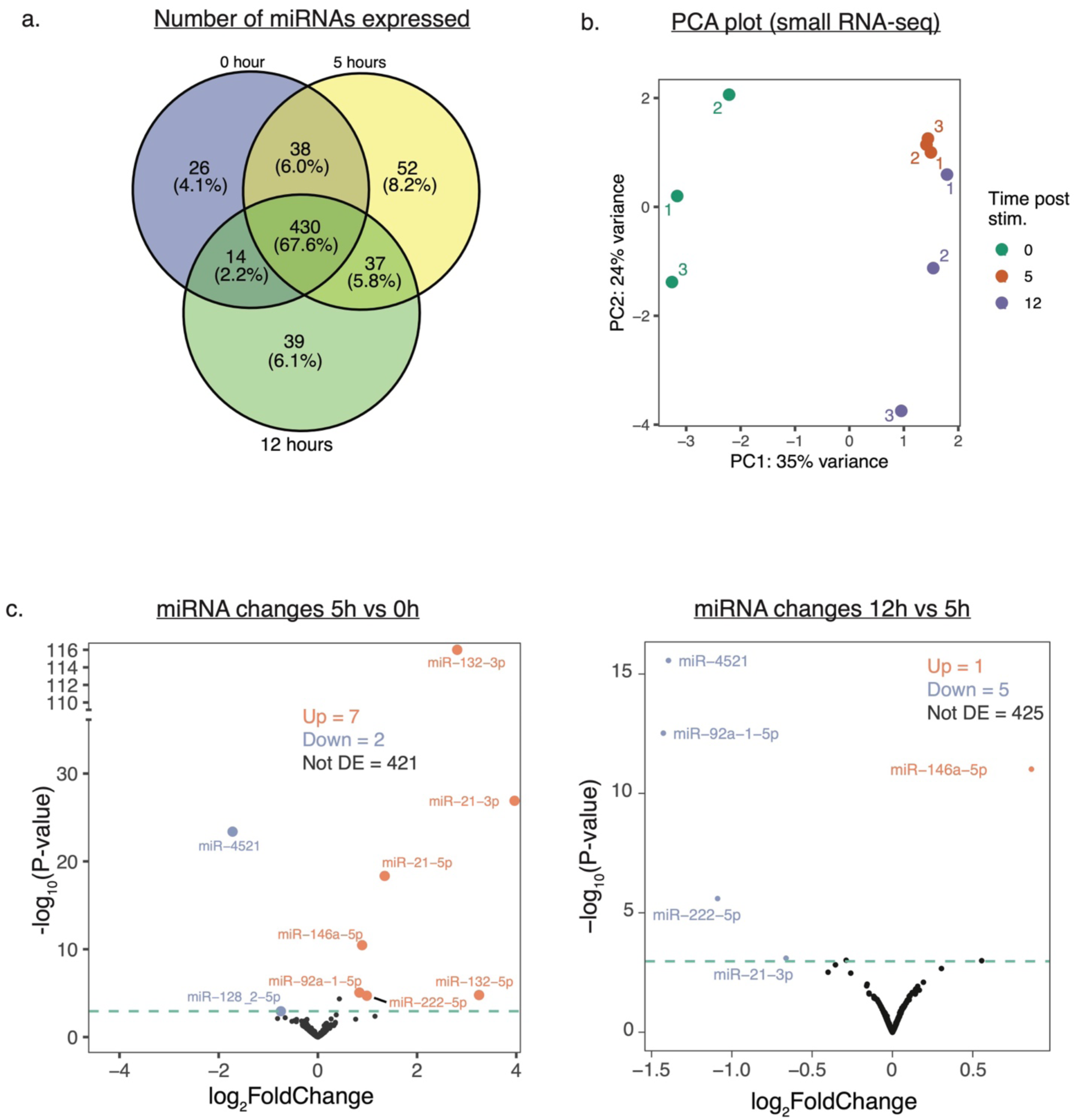
miRNA Expression Changes and Overlap Across Time Points. (a) Venn diagram illustrating the overlap in the number of expressed miRNAs across three time points (0h, 5h, and 12h). Each circle represents a different time point, with overlapping regions showing miRNAs expressed at multiple time points. (b) PCA plot of small RNA-seq data showing samples colored by the time point (0h, 5h, and 12h) post-stimulation. (c) Volcano plots showing changes in miRNA expression at 5 hours and 12 hours after activation. Each point represents an individual miRNA. The x-axis displays log2 fold changes, while the y-axis shows -log10(p-value), for significance. The threshold used was FDR (BH) ≤ 5% and |log2FC| ≥ 1.

### Identification of Validated miRNA Targets Highlights Immune Pathway Activation in T Cells

Next, we aimed to identify mRNA targets for the differentially regulated miRNAs in our data. miRNA targets can be identified by prediction tools [43–47] or from experimental methods [17,48–52]. Prediction tools commonly evaluate complementarity between miRNA seed regions and mRNA-sequences, thermodynamic stability, evolutionary conservation, and accessibility of target sites. However, these tools often have high false positive rates, as many predicted interactions may not be biologically relevant [53,54]. To improve reliability, a combination of various prediction tools and experimental methods is frequently employed, though this approach can be resource-intensive and time-consuming [55]. We therefore focused on identifying targets from databases that only report experimentally validated targets [47]. Specifically, we sourced target data from three well-established databases, miRecords [56], miRTarBase [57], and TarBase [58], each reporting only experimentally validated miRNA targets. From these databases combined, we identified approximately 17,000 experimentally validated target genes across all differentially expressed miRNAs from our dataset, with target counts for each miRNA ranging from 100 to 5,000 (Fig 6a, Suppl Table S6).

**Figure 6.**
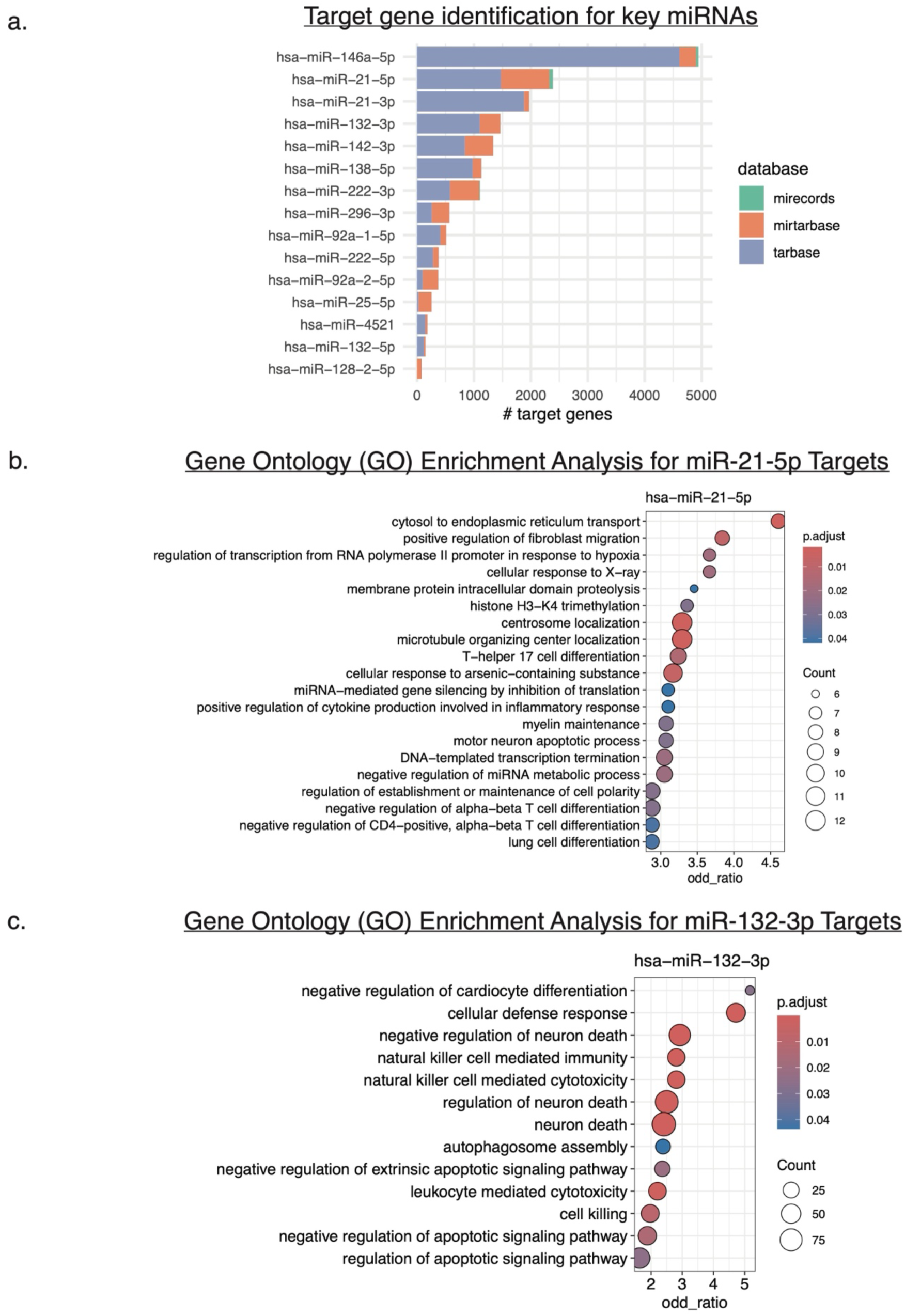
Target Gene Enrichment Analysis for Key miRNAs. (a) Bar plot displaying the number of target genes associated with each miRNA across three databases: miRecords, miRTarBase, and Tarbase. (b-c) GO term enrichment analysis for hsa-miR-21-5p (b) and hsa-miR-132-3p (c) target genes. The dot plot shows enriched biological processes, with color indicating the adjusted p-value (p.adjust) and dot size representing the gene count for each process. The threshold used was FDR (BH) ≤ 5%.

Gene Ontology analysis revealed significant enrichment in immune-related pathways within predicted mRNA targets of miRNA-21-3p, miRNA-21-5p, miRNA-128-2-5p, miRNA-222-5p, miRNA-296-3p, and miRNA-25-5p (Fig 6b-c, Suppl Fig 8-9). However, mRNAs are often regulated by multiple miRNAs binding in tandem, which can amplify the regulatory effects [59,60]. We therefore assessed how many of the predicted target genes in our dataset were regulated by multiple differentially expressed miRNAs. Of the 17,000 predicted target genes, 2416 are predicted to have 2-3 binding sites, and 463 are predicted to have 4 or more binding sites for the differentially regulated miRNAs we observe (Suppl. Table S6). Gene ontology analysis of the mRNAs with 4 or more miRNA sites revealed multiple enriched biological pathways, with translation emerging as one of the most prominent (Suppl Fig 10). This further underscores the importance of translation-specific gene regulation during early T cell activation.

### mRNAs Exhibiting Positive Regulation Are Targeted by Multiple miRNAs

We examined the effect of multiple miRNA interactions on the regulation of mRNA levels and translation. Specifically, we assessed how the predicted target genes in our dataset were regulated in the different TE categories (forwarded, buffered, exclusive, and intensified). Interestingly, mRNA targets with ≥4 predicted DE miRNA interactions showed greater positive fold changes after 5 hours of activation in the intensified set of mRNAs (Fig 7). Intensified DTEs (Differentially Translated Elements) are differentially translated genes where RPF levels increase more in the same direction as mRNA levels, thereby intensifying the change in gene expression. However by 12 hours, the intensified DTEs with ≥4 predicted DE miRNA interactions trend towards being translationally repressed (Fig. 7). By contrast, buffered and exclusive DTEs with ≥4 predicted DE miRNA interactions which were mostly repressed at 5 hours become more evenly divided between positive and negative fold changes in translation (Fig. 7). As expected, the “forwarded” category, characterized by proportional changes in both mRNA and RPF levels (log₂TE values near 0) did not exhibit clear peaks of positive or negative regulation. In summary, our findings suggest that mRNAs targeted by ≥4 DE miRNAs may result in positive translation regulation at 5 hours or 12 hours, suggesting that multiple miRNAs may contribute to enhanced gene expression during T cell activation. However further validation is needed to confirm a direct regulatory effect for the combined action of these DE miRNAs.

**Figure 7.**
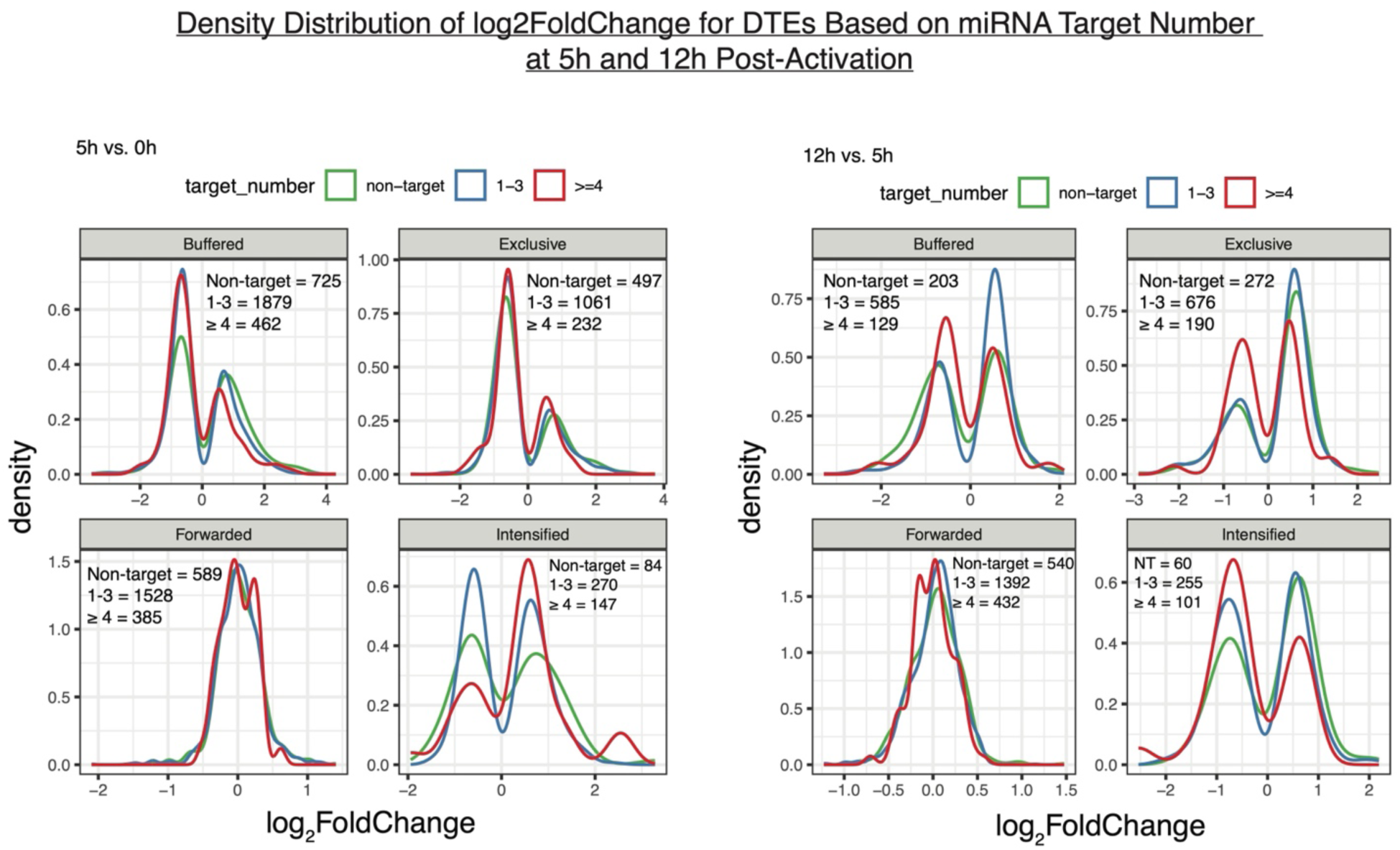
Density Distribution Analysis of Differentially Translated Elements (DTEs) Across Time Points based miRNAs targeting. Density plots categorized by regulatory patterns (Buffered, Exclusive, Forwarded, Intensified) at 5h vs 0h and 12h vs 5h. The plots compare the log2FoldChange of targets based on the number of DE miRNAs predicted to target them (0, 1-3, >=4) to illustrate the impact of miRNA targeting on gene expression changes. Note that forwarded genes cannot be classified as “upregulated” or “downregulated” based on TE, as both RNA and RPF levels change proportionally, resulting in the absence of distinct regulatory peaks.

Next, we leveraged the mRNA-seq and ribosome profiling data to investigate the differential regulation of specific target genes with ≥4 DE miRNAs during T cell activation in our data. We compared the expression profiles of mRNAs, RPFs of all miRNA targets, and the miRNA itself across three time points (0 hour, 5 hours, and 12 hours), and calculated Z-scores by subtracting the mean relative expression value across all time points from each individual expression value of each dataset (miRNA, mRNA, or RBP) and then dividing by the standard deviation. This normalization centered the data, allowing positive and negative Z-scores to reflect expression levels above or below the mean, respectively. Although differential effects are masked when using Z-scores over all genes with differential TE in a regulatory category (Suppl Fig 11-12), individual genes showed striking patterns of regulation. For example, mRNA levels for *DDX3X*, *BTG2*, *MYC*, and *AGO2* were elevated at 5 hours, concomitant with increases in miRNA levels, and then decreased at 12 hours (Fig. 8). RPFs for *DDX3X*, *BTG2*, and *MYC* remained relatively constant at 5 hours, then dropped at 12 hours in parallel with mRNA levels (Fig. 8). These results are consistent with translation repression preceding mRNA decay for these three transcripts [61] Interestingly, the expression pattern for *EIF4G3* and *PABPC1* follow a different pattern. For these genes, relative mRNA levels increase at 5 hours and remain constant at 12 hours, tracking the relative levels of the DE miRNAs (Fig. 8). By contrast, RPFs for these genes initially decrease at 5 hours, then either increase (*EIF4G3*) or stay at similar levels (*PABPC1*) (Fig. 8). Taken together, the different patterns of DE miRNA, mRNA, and RPF levels seen for these 6 genes suggest that further experiments are needed to dissect the contribution of miRNAs to their regulation at both the mRNA and translation levels during T cell activation.

**Figure 8.**
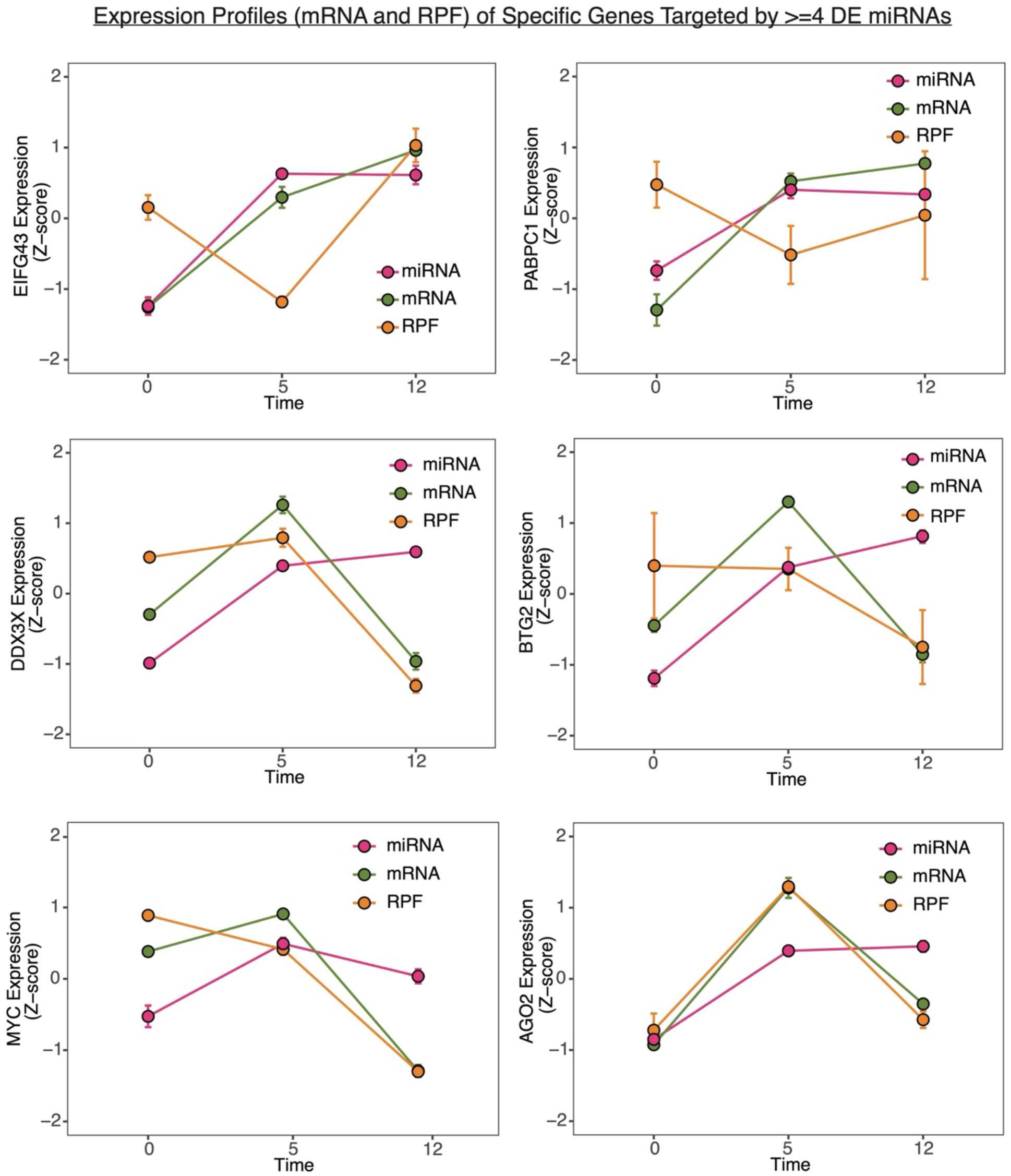
Expression Profiles (mRNA and RPF) of Selected Genes Targeted by >=4 miRNAs Across Time Points. Dot plots showing the mRNA and RPF expression (scaled to unit variance, Z-score) of selected genes, along with the average expression of DE miRNAs predicted to target them at 0h, 5h, and 12h. Error bars indicate the standard error of the mean (s.e.m.), first from biological replicates for mRNA and RPF, and then across different genes for mRNA and RPF and biological replicates for miRNA data.

## Discussion

### T Cell Activation and Gene Regulation

T cell activation triggers substantial changes in gene expression, propelling cells from a resting state to an active, proliferative, and effector state. This study assessed these changes at early (5 hours) and mid (12 hours) activation time points in Jurkat cells using mRNA-seq to examine mRNA expression, ribosome profiling for protein translation, and small RNA-seq to monitor miRNA expression. Early T cell activation is a critical phase in the immune response because it determines the magnitude, quality, and specificity of downstream immune functions. Although Jurkat cells have limitations in fully representing primary T cell activity, they are widely accepted for CD4+ T cell molecular studies [62–64] and served as our model for the present study. In our data, most gene expression alterations were identified at the 5 hours mark, with minimal additional changes by 12 hours (Fig 2-3, Suppl Fig 2b), suggesting that gene expression adjustments are primarily an early event in T cell activation. This pattern was also evident in T-cell specific genes (Suppl Fig 2c). We also observed more translational upregulation than downregulation of all genes at 5 hours, whereas transcriptional up- and downregulation were roughly equal. However, for T cell–specific genes, both transcription and translation showed more upregulation than downregulation (Suppl Fig 2c). We detected several T cell activation related genes (Fig 3a-d) in both the 5 hours and 12 hours datasets, including early activation marker *CD69*, *CD83* and *LAT*.

### Translation efficiency changes and T cell activation

We also leveraged our mRNA-seq and ribosome profiling datasets to calculate translation efficiency and further classify the translation efficiency changes into four regulatory categories [34], based on changes in mRNA and ribosome footprint levels. This classification enabled us to use GO analysis to gain insight into how distinct gene programs behave over time during T cell activation. Since proliferation and protein translation are key events in this process [2,3], we focused on how genes associated with these pathways are regulated. Our data showed that, for T cell or immune-specific genes, both mRNA and ribosome footprint levels changed to similar extents at 5 hours, indicating transcription-driven regulation. However, by 12 hours, translation emerged as the dominant regulatory layer, amplifying the transcriptional changes in immune-related pathways. By contrast, several cell division–specific gene categories followed the opposite pattern, exhibiting a translational boost at 5 hours that reverted to transcription-driven regulation by 12 hours. We also observed that pathways associated with ribosome biogenesis were translationally regulated at 5 hours, with some aspects buffered at both 5 hours and 12 hours. These dynamic patterns suggest that distinct regulatory strategies like transcriptional activation, translational amplification, and buffering are selectively deployed across gene programs. Classifying genes based on combined mRNA and ribosome footprint changes provides a deeper understanding of how key events such as translation and cell proliferation are orchestrated during T cell activation. Furthermore, measuring gene expression at the early 5- hour time point revealed regulatory changes that would have been missed by analyzing only the 12-hour time point. To our knowledge, no previous study has precisely delineated the temporal regulation of these pathways during T cell activation, which is essential for understanding the gene regulatory networks that drive immune responses.

### T Cell Activation and Micro-RNAs

We detected over 600 miRNAs across all time points during early and mid T cell activation, with 430 miRNAs consistently expressed across conditions. Principal component analysis revealed a clear separation between non-activated (0h) and activated (5h, 12h) cells, indicating activation-associated shifts in the small RNA landscape. While most miRNAs maintained stable relative expression, a small subset exhibited differential expression over time. Overall, our analysis revealed 9 miRNAs differentially expressed at 5 hours and 6 miRNAs differentially expressed at 12 hours (Fig 5c). Both the 5p and 3p strands of miRNA-132, miRNA-21 and miR-4521 showed differential expression at 5 hours and 12 hours, consistent with previous findings [17]. The functions in immune responses of some of the DE miRNAs we identified, miRNA-132, miRNA-222, and miRNA-21, are well-documented [65–70]. However, few studies have explored the specific regulation of their 5p and 3p strands in T cells [71,72].

Identifying miRNA targets ideally requires experimental, ligation-based techniques [48,73]. Due to validation challenges, computational tools are frequently employed despite high false-positive rates and assumptions about seed matching and conservation. Experimentally validated databases like miRTarBase and DIANA-TarBase, utilizing qRT-PCR and CLIP-Seq, enhance accuracy [74,75]. We identified targets of the DE miRNAs using three databases: miRTarBase, miRecords, and TarBase. These databases also provided information on experiments previously conducted to validate these targets (Suppl Table S6). Our data revealed overlapping targets among differentially expressed (DE) miRNAs (Suppl. Table S5), suggesting potential cooperative regulation. Conversely, we also observed instances of single genes being targeted by multiple DE miRNAs. Based on these observations, we next examined the regulation of genes grouped by the number of predicted miRNA interactions: 1, 2–3, and ≥4. Specifically, we aimed to understand how translation efficiency is influenced by the number of predicted miRNA interactions. Interestingly, genes for which translation intensified the transcriptional effect were more likely to be targeted by ≥4 miRNAs (Fig 7 & Suppl Fig 11-12). Note that translation can intensify both upregulation or downregulation of transcription. Therefore, examining mRNA and RPF patterns may not distinctly show simple up- or down regulation but rather reflect the average of all such effects (Suppl Fig 11-12).

The positive regulation noted in Fig 7 for genes in the intensified category is an emerging regulatory mechanism at both transcriptional and translational levels. For example, miR-373 has been shown to activate transcription by binding to complementary promoter sequences, thereby recruiting RNA polymerase [76]. At the translational level, miRNAs such as let-7 can activate translation in quiescent cells by binding AU-rich elements in the 3′ UTR, recruiting *AGO2* and *FXR1* under nutrient deprivation [25]. miRNAs have also been shown to enhance gene expression in quiescent cells, such as oocytes [77]. Additional examples of miRNA-driven gene activation include their binding to the 5′ UTR of mRNAs that code for ribosomal proteins, particularly under conditions of amino acid starvation [26]. These examples underscore the adaptability of miRNAs in gene regulation, responding flexibly to cellular conditions and needs. Interestingly, GO analysis of targets regulated by ≥4 miRNAs showed significant enrichment for genes involved in “translation,” further highlighting the importance of translational upregulation in T cell activation (Suppl Fig 10). Previous studies have demonstrated the phased roles of miRNAs, with mRNA decay typically preceding translational repression, except under specific conditions. For instance, miRNAs in embryonic stem cells can inhibit translation without degrading mRNA, with *DDX6* facilitating this process by engaging the translational machinery [78]. Deadenylation complexes such as PAN2/PAN3 and CCR4–NOT further reinforce this repression, allowing for dynamic adjustments in gene expression [79]. In zebrafish, miR-430 initially reduces ribosome occupancy on target mRNAs before promoting decay, a two-stage regulatory process that ensures rapid and sustained silencing [80]. Our findings also revealed various instances where either repression or decay could precede the other (Fig 8).

### Future Directions

Our data showed that positively regulated miRNA targets at both mRNA and RPF level were targeted by multiple miRNAs. The relationship between miRNA seed sequences and target upregulation remains poorly understood, but miRNA interactions with AU-rich elements [25,81] and motifs like the 5′TOP motif [26] are promising areas of study. It is possible that non-seed-based recognition may influence upregulated targets, warranting further investigation [81]. Research on miR-430 in zebrafish shows that miRNA-mediated translational repression predominantly affects initiation, where miR-430 decreases ribosome occupancy on target mRNAs without affecting elongation, highlighting miRNAs’ precise influence over translation initiation [61]. Further exploration of the mechanisms underlying translational up and down-regulation at the individual mRNA level could provide deeper insights into miRNA function. Our study presents a comprehensive framework for understanding gene regulation in T cells, shedding light on miRNA effects during T cell activation. Further investigation of miRNA binding elements in 5′-UTRs, along with translation initiation data from ribosome profiling of positively regulated targets, would enhance the mechanistic understanding of this regulation. Additionally, miRNA knockout studies could help determine the direct effects of miRNAs on target upregulation while ruling out indirect contributions from RNA-binding proteins.

## Conclusions

Our integrative analysis of transcriptional and translational changes during early T cell activation provides new insights into how gene expression is rapidly reprogrammed in response to stimulation. We demonstrate that the most substantial changes in both mRNA abundance and translation efficiency occur by 5 hours post-activation, highlighting the acute and coordinated nature of early T cell responses. This early wave of gene regulation prominently affects pathways related to translation, proliferation, and immune signaling.

By combining ribosome profiling with small RNA and mRNA sequencing, we systematically identified 9 miRNAs differentially expressed during early activation, most of which exhibit expression changes within the first 5 hours. Importantly, we found that 6 of these miRNAs are predicted to target genes involved in immune-related pathways. Analysis of miRNA binding site density and target gene expression revealed a nuanced regulatory landscape. Contrary to a uniform model of miRNA-mediated repression, our data suggest a context-dependent effect: genes with ≥4 predicted miRNA binding sites were often upregulated during early activation, especially within translation-intensified categories, before transitioning toward repression at later time points. These findings suggest a dynamic model in which miRNAs initially support a burst of translation for key proliferative genes, potentially by buffering expression noise or stabilizing transcripts, followed by canonical repression as the activation program progresses. This temporal regulation underscores the complexity of miRNA function in immune responses, extending beyond simple target degradation to include fine-tuning of protein synthesis in a phase-specific manner.

Overall, our study lays the groundwork for understanding how miRNAs contribute to the temporal orchestration of gene expression during T cell activation. Future experiments in primary T cells and *in vivo* models will be essential to validate specific miRNA–target interactions and to define the functional consequences of this layered regulation. Such insights could ultimately inform the development of miRNA-based strategies for modulating T cell function in immunotherapy and autoimmune disease.

## Methods

### Cell culture

Human Jurkat Clone E6-1 (ATCC TIB-152) was obtained from the Berkeley Cell Culture Facility. Cells were routinely tested for mycoplasma contamination at the UC Berkeley Cell Culture Facility. Cells were maintained in RPMI 1640 Medium (ATCC modification) with 10% FBS (VWR Seradigm Life Science) and 0.01% Penicillin-Streptomycin (10,000 U/mL) (ThermoFisher). The cells were maintained between 1 x 10^5^ cells ml^−1^ to 1 × 10^6^ cells ml^−1^.

### Cell stimulation

Jurkat cells were activated using anti-CD3/anti-CD28 antibodies (Tonbo) for stimulation. Flat bottom plates were coated with anti-CD3 antibody at a 10 μg/mL concentration at 4 °C for at least 12 hours, and anti-CD28 antibody was added to the cell culture media at a concentration of 5 µg/mL and cells were incubated at 37 °C for the required timepoints.

### Flow cytometry and cell sorting

Flow cytometric analysis was conducted using an Attune NxT Acoustic Focusing Cytometer (ThermoFisher). For surface staining and cell sorting, cells were pelleted and resuspended in 50 μl of FACS buffer (2% FBS in PBS) containing antibodies at a 1:100 dilution (BioLegend) and incubated on ice in the dark for 30 minutes. Following staining, cells were washed twice in FACS buffer, resuspended, and analyzed. To assess cell surface expression of CD25 and CD69, gating for CD25+CD69+ cells was established based on CD25 and CD69 levels in non-activated T cells. All flow cytometry data were processed using FlowJo software (BD Life Sciences)[82].

### Transcriptome and small-RNA profiling

RNA samples were extracted from non-activated Jurkat cells or Jurkat cells activated for 5 hr and 12 hr with anti-CD3/anti-CD28 antibodies (Tonbo), using the Direct-zol RNA Miniprep kit (Zymo Research). RNA integrity was assessed using Agilent 2100 Bioanalyzer at the QB3 Genomics Facility, with RNA integrity numbers (RIN) displayed for each sample (Supp Fig 1a). RNA integrity was confirmed by RIN ranging from 8 to 9.4 (Suppl Fig 1a). Both small-RNA seq and mRNA-seq libraries were prepared from the same total RNA samples. RealSeq-AC (Somagenics) library preparation kit was used for small RNA-seq and KAPA mRNA HyperPrep kit (Roche) was used for mRNA-seq following the manufacturer’s instructions, with three biological replicates. For mRNA-seq, we mapped 28 to 35 million paired-end polyadenylated [poly(A)+] selected reads per sample.

### Ribosome profiling

Ribosome profiling was carried out with non-activated Jurkat cells or Jurkat cells activated for 5 hr and 12 hr with anti-CD3/anti-CD28 antibodies (Tonbo). 100 µg/mL cycloheximide (Sigma Aldrich) was added to the 1X Phosphate buffered saline (PBS) and the lysis buffer (20mM Tris pH 7.5, 150 mM NaCl, 5 mM MgCl_2_, 1 mM DTT) during ribosome profiling to stabilize the translatome without affecting the outcome [83]. 20 million cells were used per condition per replicate. Cell lysates were snap-frozen in liquid nitrogen before long-term storage at -80 °C or used immediately after. Cell lysates were digested with 20 U/μg of P1 nuclease at 37 °C for 60 minutes. Digested lysates were pelleted on a sucrose cushion. The P1 RPFs were size-selected from 30 – 40 nt after resolving on denaturing 15% urea-PAGE. Library preparation was carried out by OTTR library generation method [33,84–86]. OTTR reaction products were size selected in 8% Urea-PAGE gel and cDNAs were quantified by real-time PCR in order to determine the number of PCR cycles needed for library amplification [87]. Libraries were generated using standard Illumina primers. We processed 52 to 192 million single end reads per sample.

### Small RNA-Seq bioinformatics

Raw demultiplexed FASTQs were subjected to quality control using cutadapt v.1.16 [88] similar to the above. Briefly, the first base at 5’-end was trimmed (-u 1), followed by adapter removal (- a TGGAATTCTCGGGTGCCAAGG), and length filtering retaining only reads 15 nucleotides-longer (-m 15). Then, reads were aligned sequentially first to the tRNA references, tRAX reference, and mtDNA using Bowtie 2. Reads not mapping to these references were then mapped against the human miRNA hairpin reference with non-stringent settings (bowtie –norc -p 4 -m 50 -v 3). Hairpin-mapped reads were then aligned with mature miRNA reference sequences with a stricter setting (first with zero mismatches, followed by up to one mismatch). Then, each of the non-mapping reads are realigned after trimming the first base at 5’ with “*bowtie --norc -m 50 -v 0 -5 1*”. Reads not mapping after these steps were trimmed for polyA[100] and polyT[100] using cutadapt and then mapped again to mature refs (bowtie --norc -m 50 -v 1 -5 1). Reads not mapped to anything were then used as input in mirDeep2 mapper.pl and quantifier.pl routines. miRNA counts from mirDeep2 were filtered to remove miRNAs that had zero counts across any sample before proceeding. DESEq2 v. 1.42.0 was used for normalization and differential gene expression analysis [89]. Significantly differentially expressed miRNAs for each timepoint were determined based on their fold-change and an adjusted p-value < 0.05. For seed sequence analysis, we used the scanMiR v.1.6.0 with the defaultOnly = TRUE argument. Briefly, the longest transcript for each gene targeted by one of the miRNAs (as per experimental evidence in public databases) were extracted using TxDb.Hsapiens.UCSC.hg38.knownGene v.3.17.0 and BSgenome.Hsapiens.UCSC.hg38 v.1.4.5 and used with the miRNA sequence in the findSeedMatches function. Then, seed categories were collapsed into either 6mer, 7mer, 8mer, combinations of thereof, and none.

### mRNA-Seq bioinformatics

Raw demultiplexed FASTQs were subjected to quality control using cutadapt v.1.16 [88]. Briefly, the reads were trimmed at the 3’ with a quality threshold of 10 (-q 10), adapter-trimmed (-aA AGATCGGAAGAG), and then filtered out if the length was shorter than 20 nucleotides. Reads were aligned against the Gencode v.37 (GRCh38.p13) protein-coding transcripts using Bowtie v.1.0.0 [90]. Only reads aligned uniquely (-a -m 200) were kept. Additional parameters used were ‘-e 99999999 --chunkmbs 800 -I 99 -X 999 --allow-contain’. Samtools v.1.7 was used to convert SAM to BAM while excluding unmapped reads. Next, BAM files were prepared for gene expression quantification using the RSEM v.1.3 routines ‘convert-sam-for-rsem’ and ‘rsem-calculate-expression’ [91]. DESEq2 v. 1.42.0 was used for normalization and differential gene expression analysis [89]. Only genes with more than zero counts across all samples were analyzed. Log2 fold-changes were shrunken using the apeglm v1.22.1[92] or ashr v2.2-63 [93] packages. We then generated deep sequencing libraries and mapped 28 to 35 million paired-end polyadenylated [poly(A)+] selected reads per sample.

### Ribosome profiling computational analysis

The raw reads were initially trimmed using cutadapt v3.2. Briefly, a custom script was used to partition reads based on the ligated adaptor’s linker sequence. The procedure removed the adapter GATCGGAAGAGCACACGTCTGAACTCCAGTCAC followed by annotating the UMI and primer +1 base (+1T or +1C) in the FASTQ ID: “cutadapt -m 2 -a GATCGGAAGAGCACACGTCTGAACTCCAGTCAC $tmpvar.fastq | cutadapt -u 7 --

rename=’{id} NTA={cut_prefix}’ - | cutadapt -u -1 --

rename=’{id}_{comment}_TPRT={cut_suffix}’ - | cutadapt -m 25 -q 10 -”

Next, the reads were filtered with a sequential mapping scheme. The trimmed reads were mapped with bowtie initially to the rRNA reference (bowtie -m 50 --best), the tRNA reference from tRAX (-k 100 --very-sensitive --ignore-quals --np 5 --reorder), the mtDNA reference (-m 50 -v 2 --best ), ncRNA reference (-v 3 --norc -m 300 -y). Finally, reads not mapping to any of these were mapped against the human Gencode v37 GRCh38.p13 coding reference sequences (bowtie -v 2 --norc -m 250 -a -y --all). To quantify gene-level CDS occupancy, bedtools intersect (v.2.25.0) was used to remove reads that aligned outside of the +15th to - 10th codon of the CDS. Then, reads were counted using RSEM (rsem-calculate-expression-alignments -strandedness forward -seed-length 20 -sampling-for-bam), followed by filtering for genes with more than zero counts across all samples. Then, count matrix was normalized and further analyzed with DESeq2 v1.4.2. Differential translation analysis was done by fitting gene-wise negative binomial generalized linear models using DESeq2 with concatenated counts from the RNA and ribosome profiling (combined_counts ∼ hour + hour:type). Log2 fold-changes were shrunken using the apeglm [92] or ashr [93] packages. The differential translation efficiency was obtained by extracting interaction coefficients from the glm model and categorized according to the DeltaTE method [34]. Genes categorized in forwarded, exclusive, buffered, and intensified were used as input in over-representational analysis using the Gene Ontology Biological Process with clusterProfilerr v. 4.8.3 [94,95]. For the background list, all genes were used.

### miRNA target identification

To identify the targets of miRNAs, R package multiMiR v.1.22.0 [96] was used. Only the validated target genes were retrieved from miRecords, miRTarBase, and Tarbase. Over-representation enrichment analyses were performed on the target gene lists using the EnrichGO function from clusterProfiler v.4.8.3 [94,95] with minGSSize = 10 and the universe set as the union of the all targets identified. Gene ontologies with FDR ≤ 0.1 (Benjamini-Hochberg adjustment) were considered significantly enriched among the target genes for a given miRNA. miRNA seed analysis was carried out by first extracting the 3’-UTR sequences from transcripts using the package ensembldb v.2.24.0 [97] to query the databases TxDb.Hsapiens.UCSC.hg38.knownGene and BSgenome.Hsapiens.UCSC.hg38. Then, scanMiR v.1.6.0 [98] was used to find canonical miRNA seeds using the findSeedMatches function. All graphical exploration was done using ggplot2 v.3.4.4 to investigate the association between miRNA expression and its targets at the transcription and translation efficiency levels.

### Data Availability Statement

Data underlying this article are publicly available. mRNAseq, small RNA-seq and Ribosome profiling data are deposited under GEO GSE254231. The code for data processing is available upon request.

## Funding

This work was also supported by the National Institutes of Health (NIH) grants R01-GM065050 and R35-GM148352 (to J.H.D.C), and R01-GM139008 (to Nicholas T. Ingolia). L.F. was supported by a Bakar Fellows Program Award and National Institutes of Health (NIH) DP1 HL156819 (to Kathy Collins). TLC was supported by a fellowship from the Pioneer Science Initiative and D’Or Institute for Research and Education.

## Competing interests

L.F. is a named inventor on patent applications filed by the University of California describing biochemical activities of RTs used for OTTR.

## Contributions

P.M. and J.H.D.C. designed the study; P.M. performed all the molecular lab work. L.F. helped P.M. with Ribosome Profiling. T.L.C. and L.F. analyzed data with contribution from P.M. P.M. and T.C.L. contributed to the text and figure panels. P.M., T.C.L. and J.H.C.D wrote the manuscript. All authors gave final approval for publication.

## Acknowledgements

We thank the members of the J.H.D.C. laboratory for the helpful discussion and Dr. Yumi Koga and Cynthia Hermosillo for the critical reading of the manuscript. We thank K. Collins and members of the Collins lab for OTTR library generation reagents. We also thank QB3 Genomics Sequencing facility for help with library generation and sequencing for mRNAseq and small RNA-seq and for sequencing of Ribosome Profiling samples. Fig 1a was created in https://BioRender.com.

