## Supplementary Figures for "miRNA-Mediated Regulation of Gene Expression During Early Activation in Jurkat Cells"

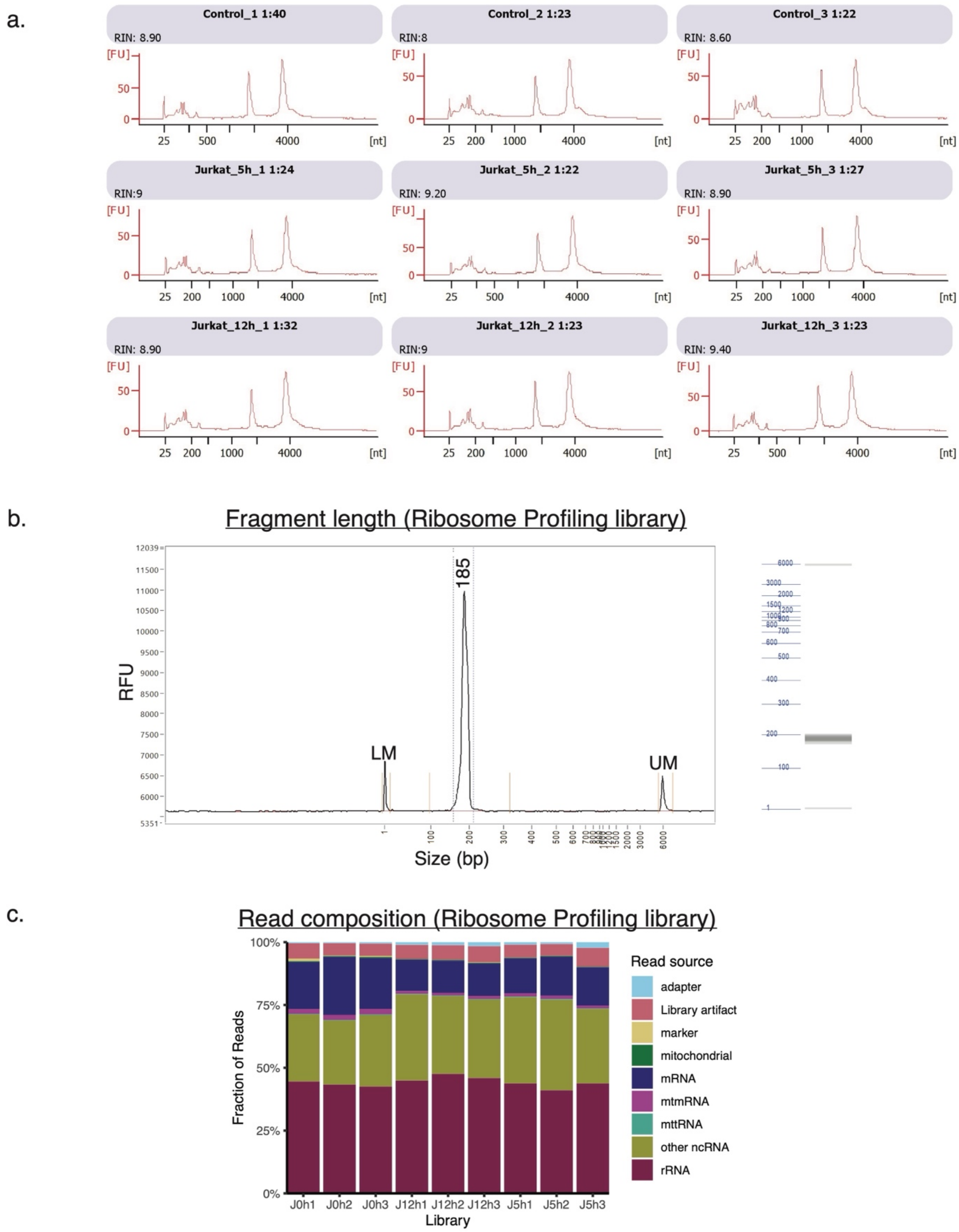


**Supplementary Figure 1. mRNA-Seq and Ribosome Profiling Data Overview.** (a) Bioanalyzer traces of RNA integrity for Jurkat cell samples at 0h (control), 5h, and 12h time points. Each plot shows the electropherogram for RNA samples, with peaks representing RNA size and quality. (b) Fragment length distribution for the ribosome profiling library with expected fragment length of 185bp for ribosome footprints. (c) Stacked bar chart illustrating the read composition of the ribosome profiling library across different libraries (samples). Colors represent various read sources, including adapters, library artifacts, mitochondrial RNA, mRNA, rRNA, and other RNA types, showing the proportion of each component.

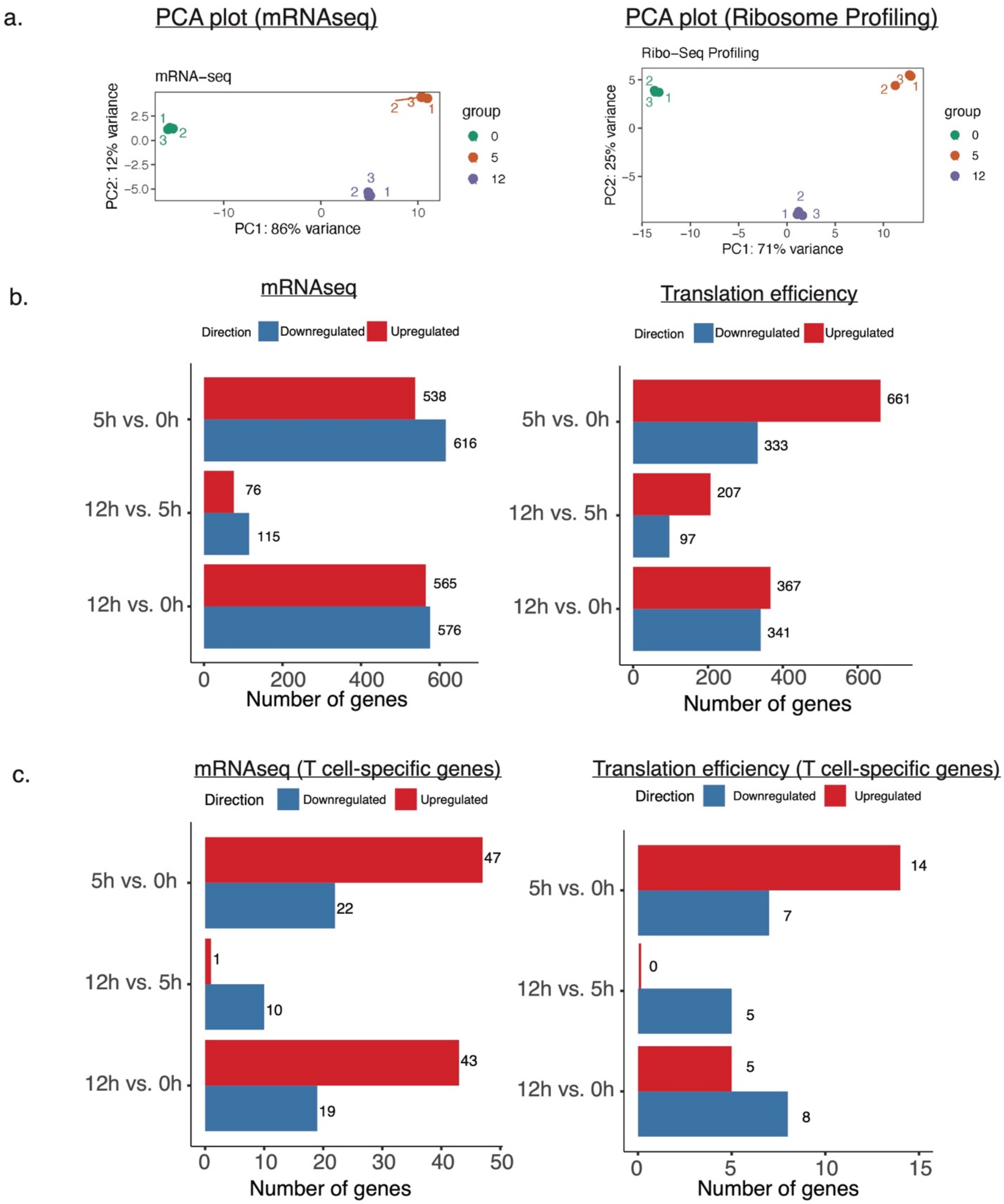


**Supplementary Figure 2. Dynamics in RNA expression and translation efficiency across time points.** (a) Principal Component Analysis (PCA) plots for mRNA-seq and ribosome profiling data. The plots show sample clustering based on variance across groups (0h, 5h, and 12h), with each group distinctly separated, illustrating the variance explained by the first two principal components. (b) Bar charts showing the number of differentially expressed genes (mRNA-seq) and differentially translated genes across time points. (c) Similar bar charts as in (b), restricted to T cell–specific genes. Bars represent counts of upregulated (red) and downregulated (blue) genes in each comparison.

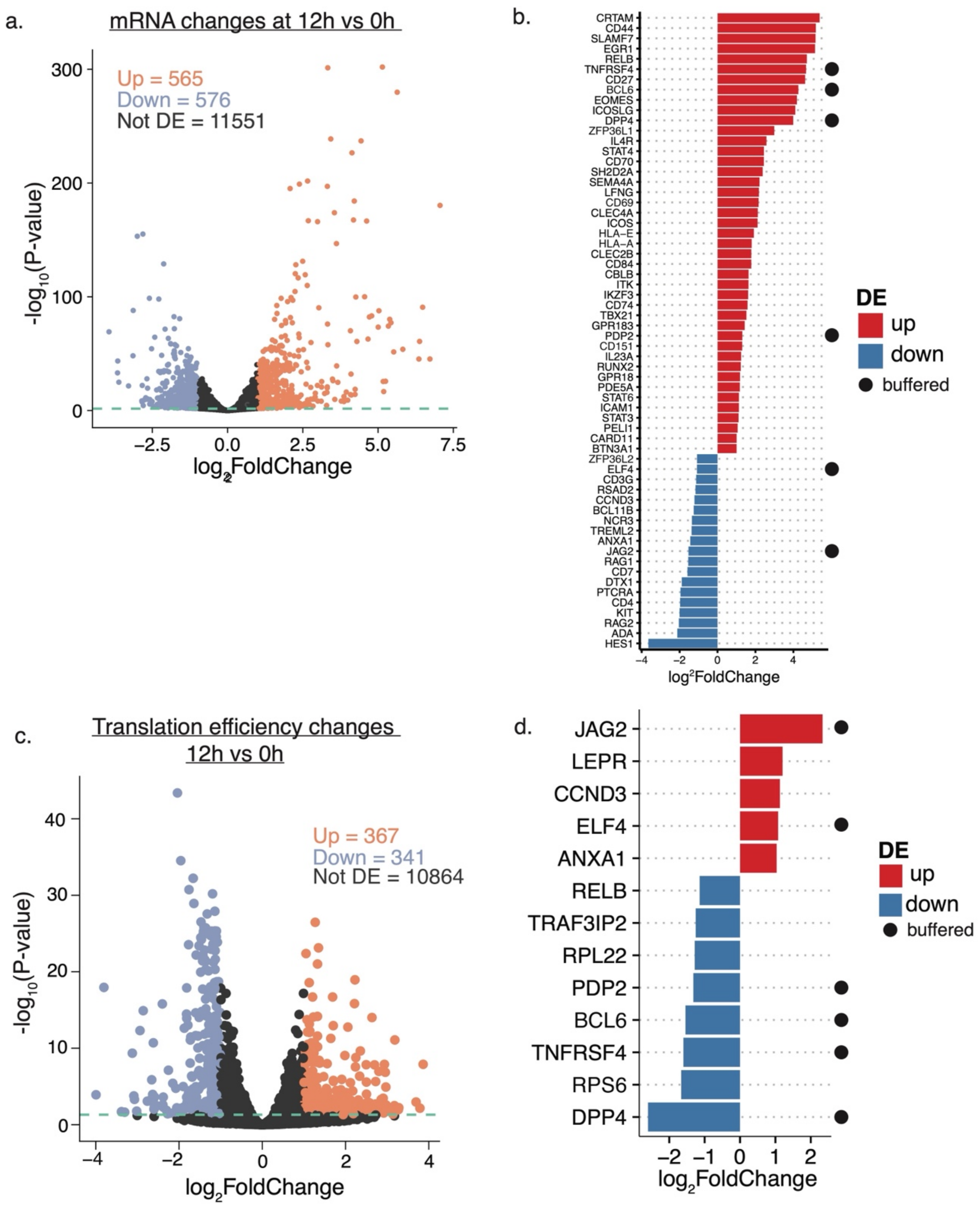


**Supplementary Figure 3. mRNA and Translation Efficiency Changes at 12 hours.** (a) Volcano plot depicting mRNA expression changes at 12 hours. (b) Bar plots showing upregulated and downregulated T cell activation-specific genes at the mRNA level at 12 hours post-activation. (c) Volcano plot depicting translation efficiency changes at 12 hours. (d) Bar plots showing upregulated and downregulated T cell activation-specific genes at the translation efficiency level at 12 hours post-activation. For volcano plots (a, c) Each dot represents a gene, with the x-axis showing log2 fold changes and the y-axis displaying -log10 p-values. Genes with significant upregulation are colored orange, and downregulated genes are in blue. For bar plots, the x-axis indicates gene names; the y-axis represents log2 fold changes. Circles indicate genes classified as buffered, as defined in Supplementary Figure 3.

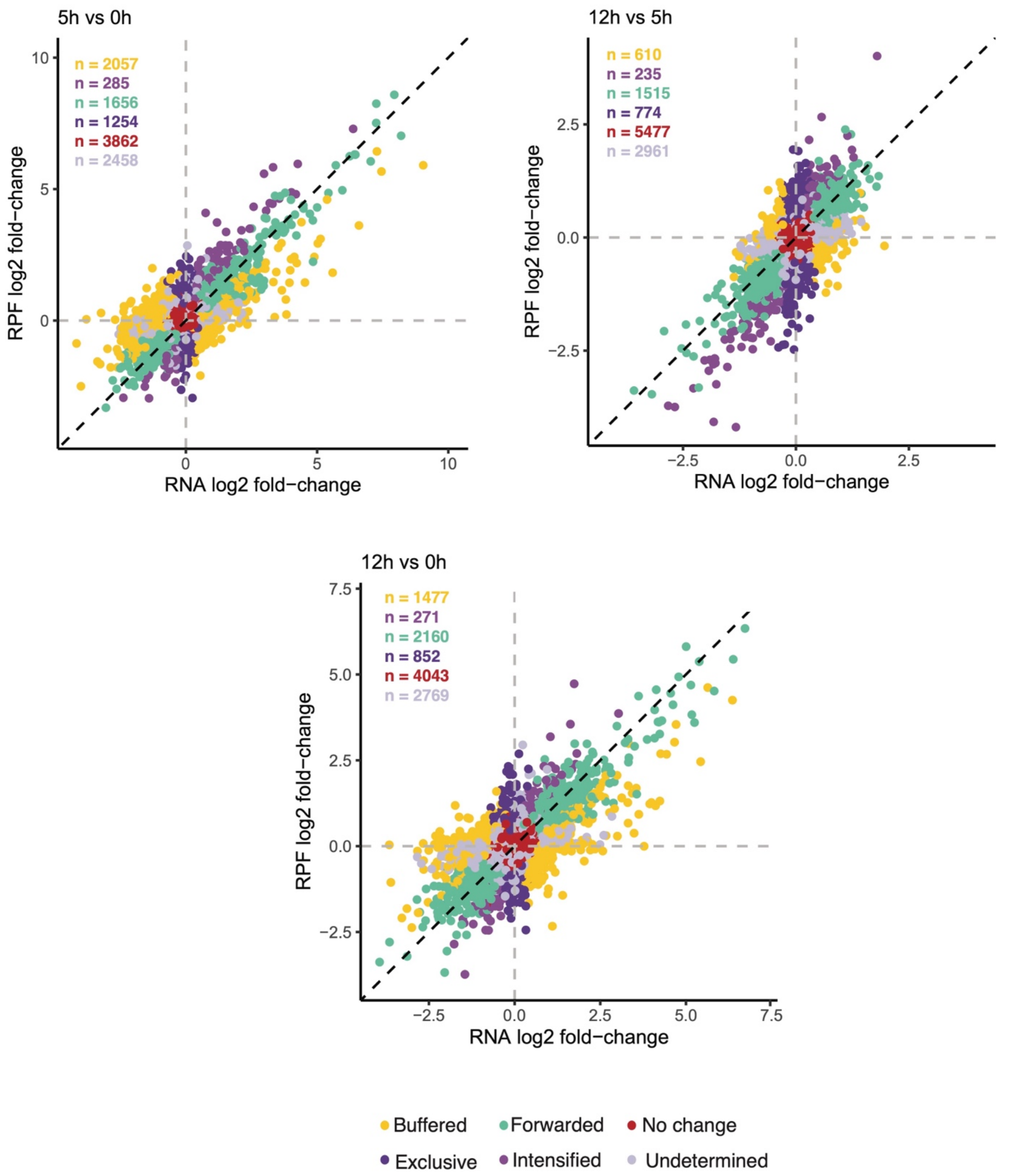


**Supplementary Figure 4. Translation Efficiency Changes at Different Time Points.** Scatter plots showing translation efficiency changes at 5h vs. 0h, 12h vs. 5h and 12h vs 0h. The plots depict ribosome profiling (RPF) log2 fold changes against RNA log2 fold changes for each comparison. Genes are categorized by regulation type: Buffered, Forwarded, Exclusive, Intensified, No Change and Undetermined. Dashed lines represent the thresholds for significant fold changes.

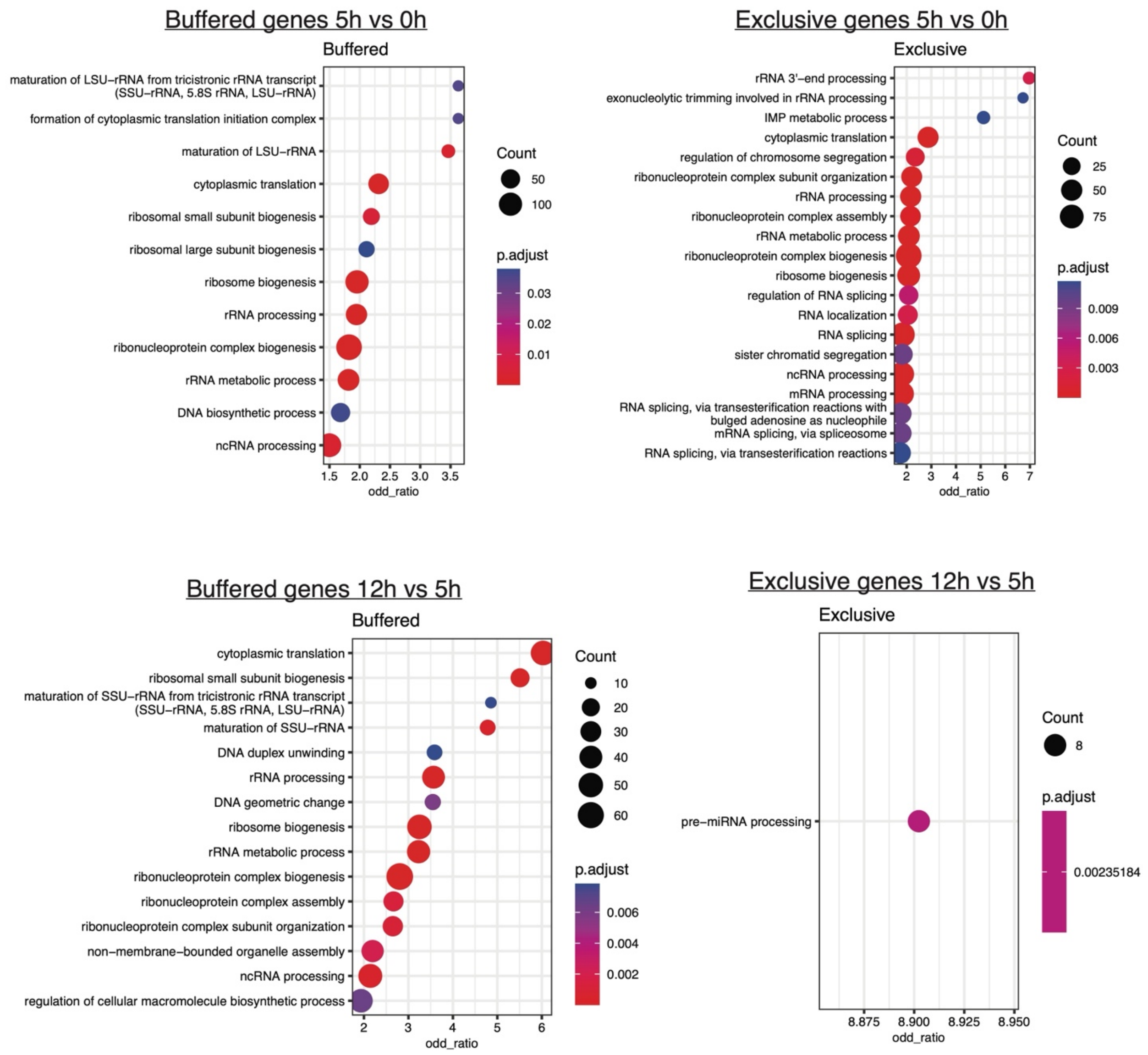


**Supplementary Figure 5. GO Enrichment Analysis for Buffered and Exclusive Genes at 5h and 12h.** (Top Row): GO term enrichment for buffered (left) and exclusive genes (right) at 5h. (Bottom Row): GO term enrichment for buffered (left) and exclusive genes (right) at 12h. Each dot plot shows enriched biological processes, with the color scale representing the adjusted p-value (p.adjust) and dot size indicating the number of genes (Count) associated with each term. The threshold used was FDR (BH) ≤ 5%.

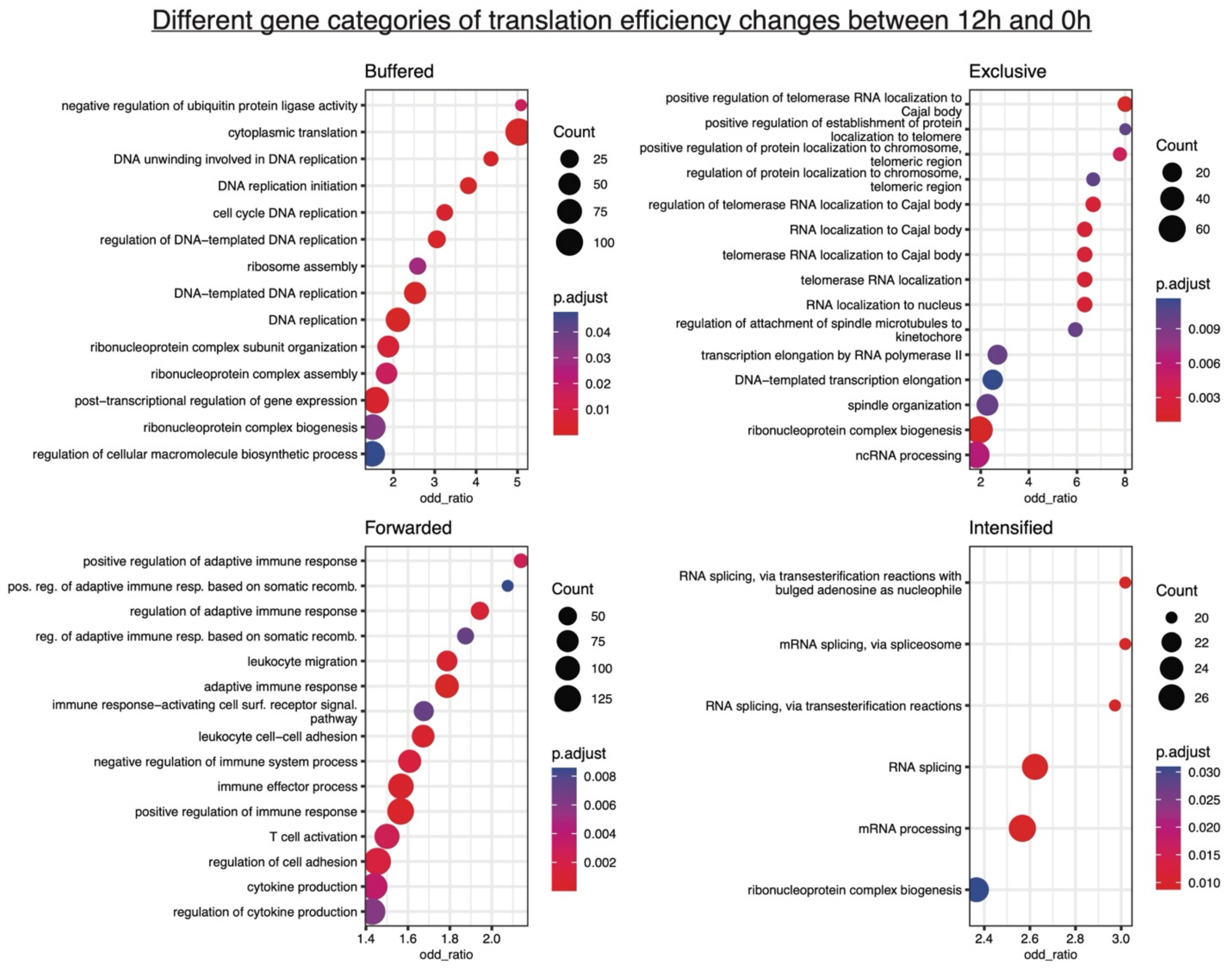


**Supplementary Figure 6. GO Enrichment Analysis for different categories of TE Genes at 12h vs 0h time point.** (Top Row): GO term enrichment for buffered (left) and exclusive genes (right) at 12h. (Bottom Row): GO term enrichment for forwarded (left) and intensified genes (right) at 12h. Each dot plot shows enriched biological processes, with the color scale representing the adjusted p-value (p.adjust) and dot size indicating the number of genes (Count) associated with each term.

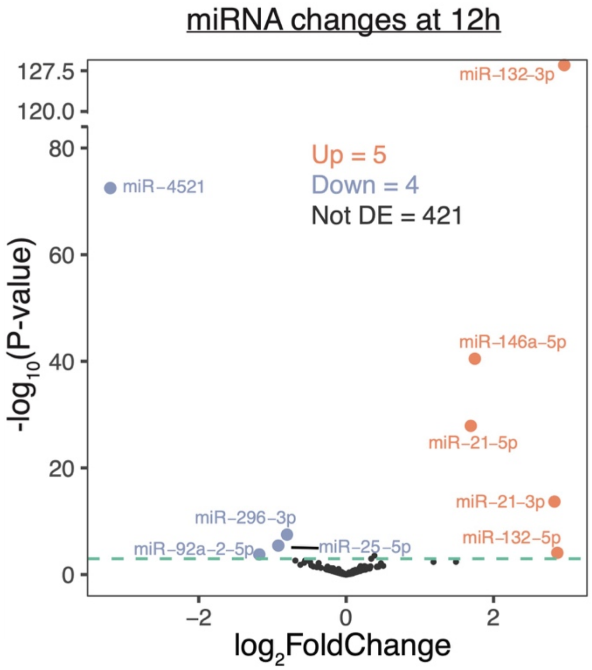


**Supplementary Figure 7.** Volcano plot showing changes in miRNA expression at 12 hours after activation. Each point represents an individual miRNA. The x-axis displays log2 fold changes, while the y-axis shows -log10(p-value), indicating significance. The threshold used was FDR (BH) ≤ 5% and |log2FC| ≥ 0.58 (~1.5 fold-change).

### Gene ontology analysis of miR-222-5p targets

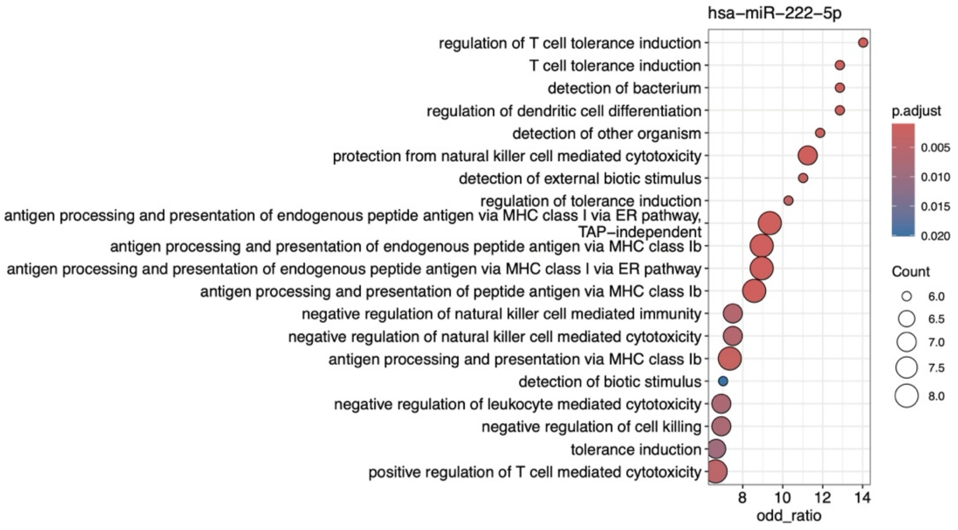


### Gene ontology analysis of miR-296-3p targets

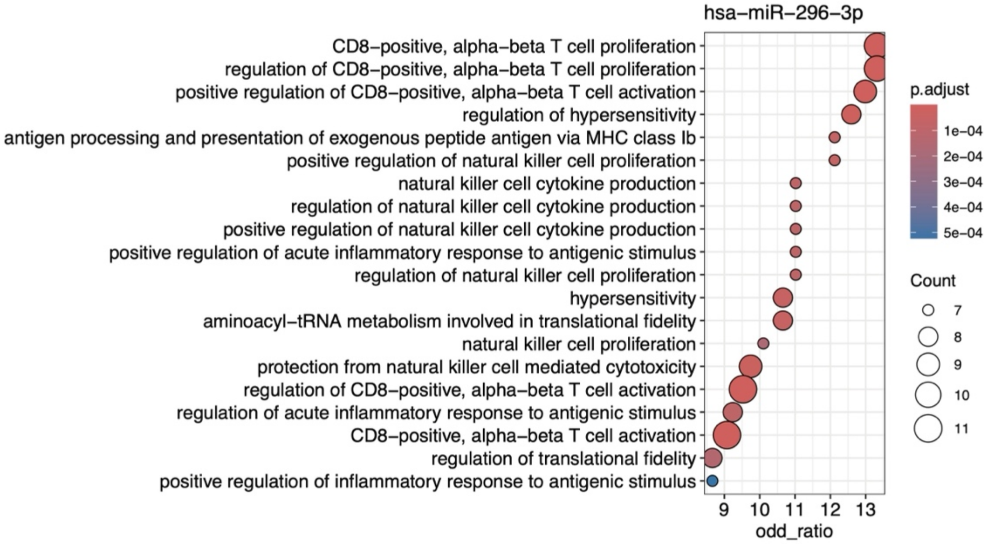


### Gene ontology analysis of miR-25-5ptargets

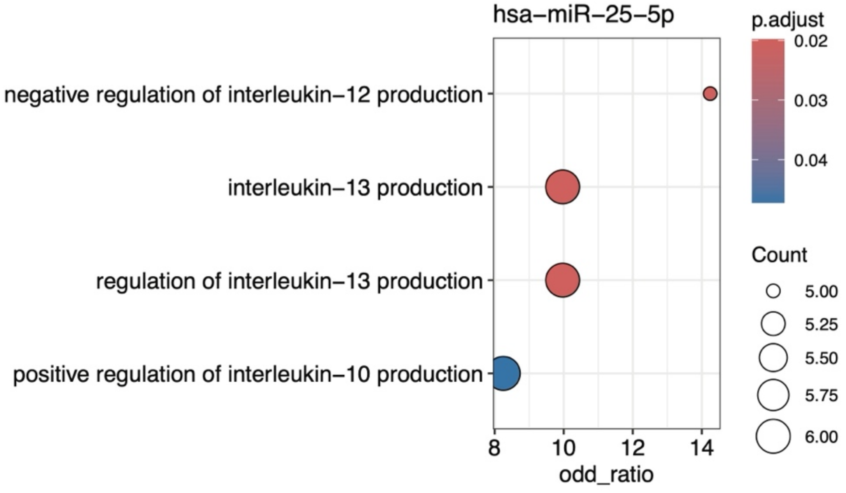

**Supplementary Figure 8. Gene Ontology (GO) Enrichment Analysis of Target Genes for miR-222-5p, miR-296-3p, and miR-25-5p.** GO enrichment analysis of target genes for selected miRNAs, illustrating the biological processes predicted to be regulated by each miRNA. Each dot plot displays enriched processes, with the color scale representing adjusted p-values (p.adjust) and dot size indicating the number of genes (Count) involved in each process.The threshold used was FDR (BH) ≤ 5%.

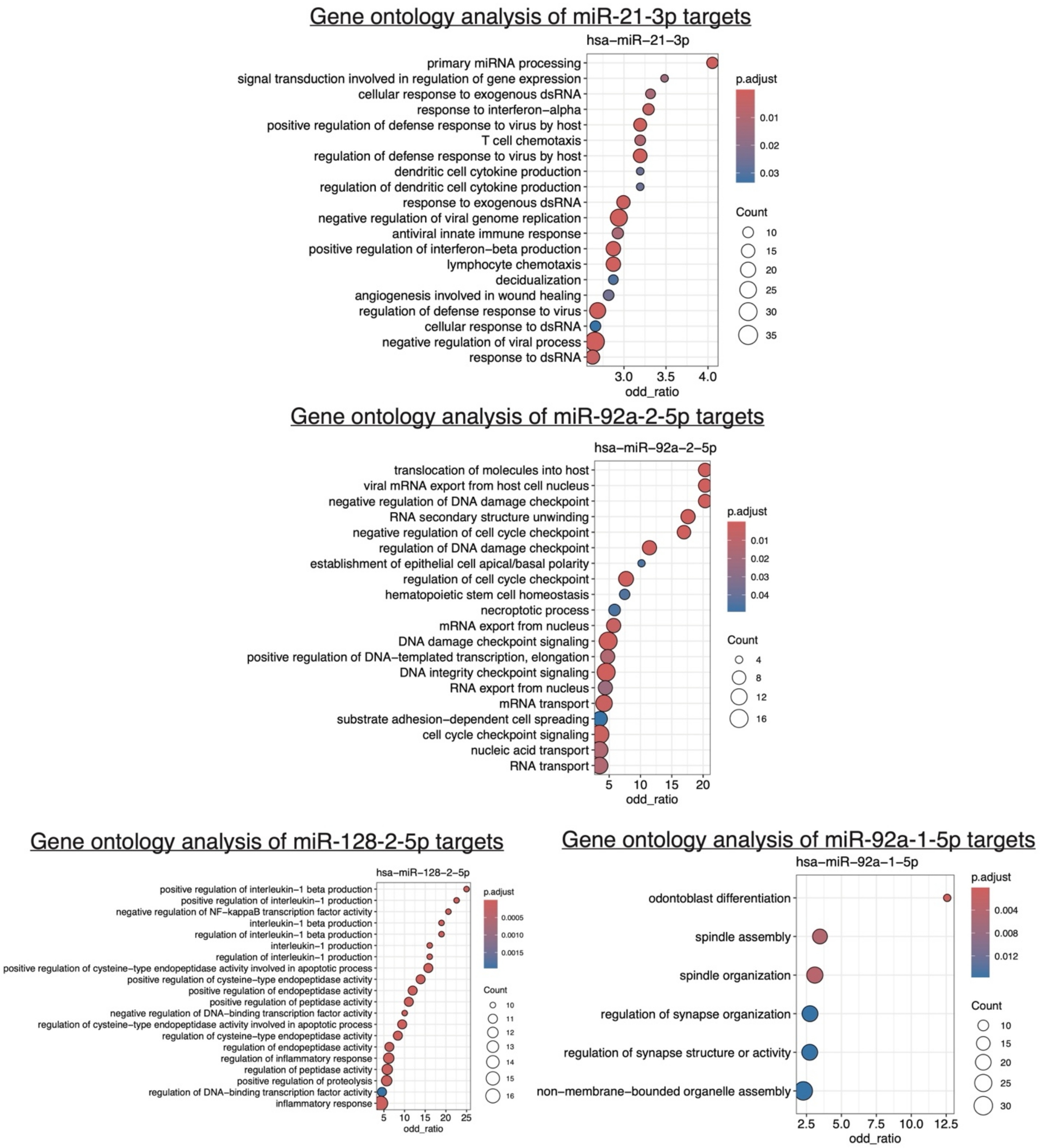


**Supplementary Figure 9. Gene Ontology (GO) Enrichment Analysis of Target Genes for miR-21-3p, miR-92a-2-5p, miR-128-2-5p and miR-92a-1-5p.** GO enrichment analysis of target genes for selected miRNAs, illustrating the biological processes predicted to be regulated by each miRNA. Each dot plot displays enriched processes, with the color scale representing adjusted p-values (p.adjust) and dot size indicating the number of genes (Count) involved in each process. The threshold used was FDR (BH) ≤ 5%.

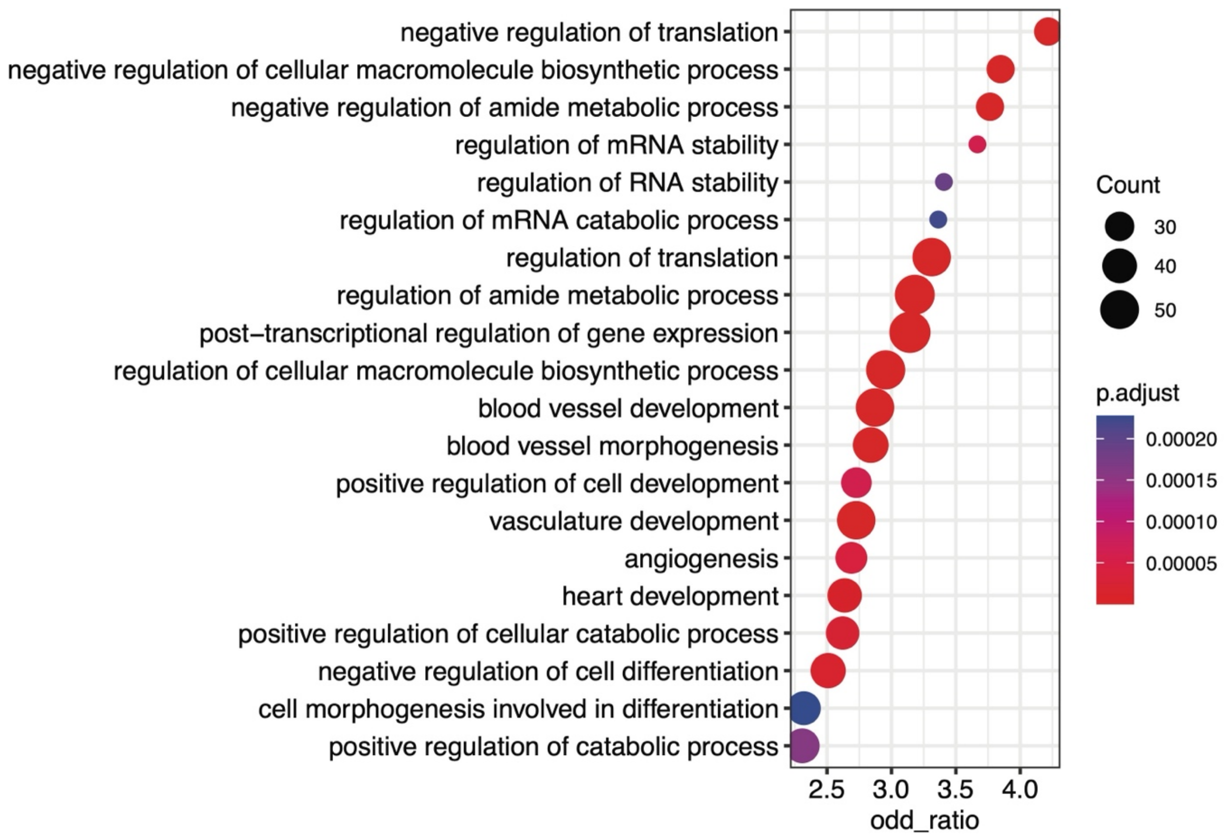


**Supplementary Figure 10. GO Enrichment Analysis of Predicted mRNA Targets Regulated by 4 or More miRNAs.** GO term enrichment for targets of 4+ miRNAs Each dot plot displays enriched biological processes, where the color scale represents the adjusted p-value (p.adjust), and the dot size indicates the number of genes (Count) associated with each term. The threshold used was FDR (BH) ≤ 5%.

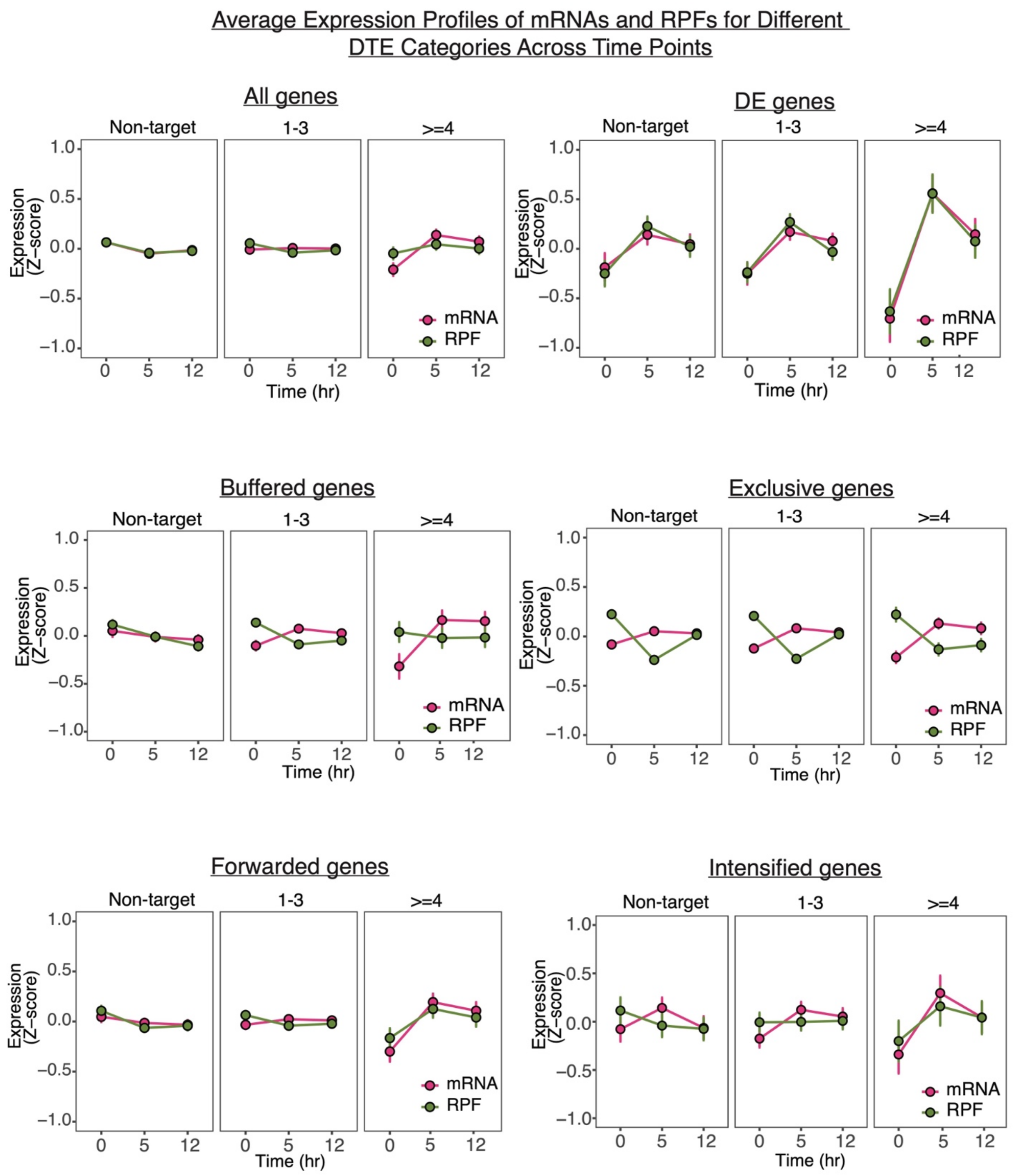


**Supplementary Figure 11. Expression Profiles of Predicted Target Gene Groups from Different Regulatory Categories Across Time Points.** Line plots showing the average expression (Z-score) of predicted target genes in specific DTE (Differential Translational Efficiency) categories. Plots include corresponding mRNA levels (pink) and ribosome-protected fragments (RPFs, green) at 0, 5, and 12 hours. Error bars indicate the standard error of the mean (s.e.m.) across biological replicates, and then calculated for the average expression of all target genes, for both mRNA and RPF. The threshold used was FDR (BH) ≤ 5%.

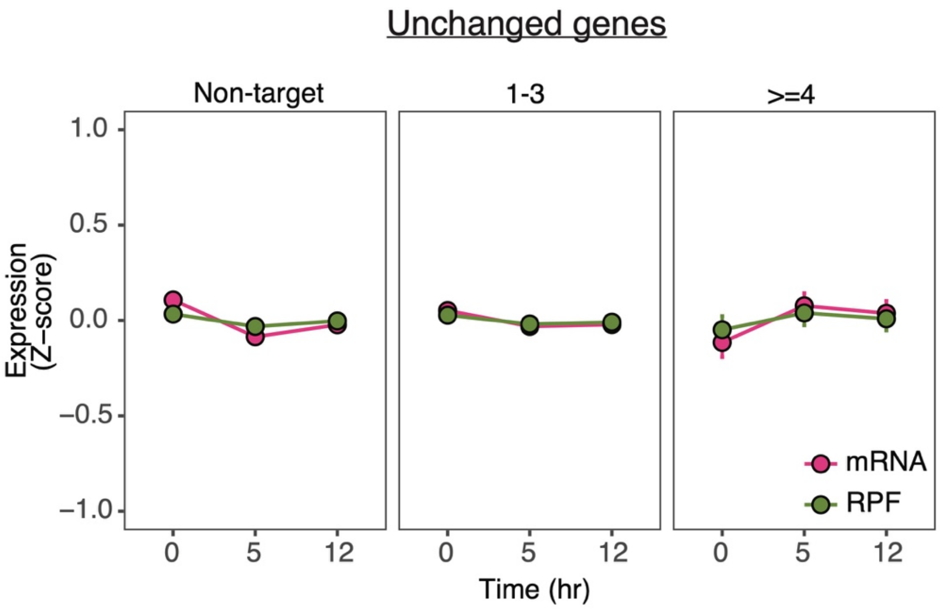


**Supplementary Figure 12. Expression Profiles of Predicted Target Genes from "Unchanged genes" Category Across Time Points.** Line plots showing the average expression (Z-score) of target genes in the Unchanged gene category. Plot includes corresponding mRNA levels (pink) and ribosome-protected fragments (RPFs, green) at 0, 5, and 12 hours. Error bars indicate the standard error of the mean (s.e.m.) across biological replicates, and then calculated for the average expression of all target genes, for both mRNA and RPF. The threshold used was FDR (BH) ≤ 5%.
